# SARS-CoV-2 spike S375F mutation characterizes the Omicron BA.1 variant

**DOI:** 10.1101/2022.04.03.486864

**Authors:** Izumi Kimura, Daichi Yamasoba, Hesham Nasser, Jiri Zahradnik, Yusuke Kosugi, Jiaqi Wu, Kayoko Nagata, Keiya Uriu, Yuri L Tanaka, Jumpei Ito, Ryo Shimizu, Toong Seng Tan, Erika P Butlertanaka, Hiroyuki Asakura, Kenji Sadamasu, Kazuhisa Yoshimura, Takamasa Ueno, Akifumi Takaori-Kondo, Gideon Schreiber, The Genotype to Phenotype Japan (G2P-Japan) Consortium, Mako Toyoda, Kotaro Shirakawa, Takashi Irie, Akatsuki Saito, So Nakagawa, Terumasa Ikeda, Kei Sato

## Abstract

Recent studies have revealed the unique virological characteristics of Omicron, the newest SARS-CoV-2 variant of concern, such as pronounced resistance to vaccine-induced neutralizing antibodies, less efficient cleavage of the spike protein, and poor fusogenicity. However, it remains unclear which mutation(s) in the spike protein determine the virological characteristics of Omicron. Here, we show that the representative characteristics of the Omicron spike are determined by its receptor-binding domain. Interestingly, the molecular phylogenetic analysis revealed that the acquisition of the spike S375F mutation was closely associated with the explosive spread of Omicron in the human population. We further elucidate that the F375 residue forms an interprotomer pi-pi interaction with the H505 residue in another protomer in the spike trimer, which confers the attenuated spike cleavage efficiency and fusogenicity of Omicron. Our data shed light on the evolutionary events underlying Omicron emergence at the molecular level.

**Highlights:**

- Omicron spike receptor binding domain determines virological characteristics
- Spike S375F mutation results in the poor spike cleavage and fusogenicity in Omicron
- Acquisition of the spike S375F mutation triggered the explosive spread of Omicron
- F375-H505-mediated π-π interaction in the spike determines the phenotype of Omicron

## Introduction

Since the emergence of SARS-CoV-2 at the end of 2019, this virus has diversified spectacularly. In April 2022, the WHO defined two variants of concern, Delta (B.1.617.2 and AY lineages) and Omicron (originally the B.1.1.529 lineage, then reclassified into BA lineages) (WHO, 2022); and currently, Omicron is the predominant variant spreading worldwide.

Even before the emergence of the Omicron B.1.1.529 lineage at the end of November 2021 in South Africa (National Institute for Communicable Diseases, 2021), SARS-CoV-2 was highly diversified from the original linage, the B lineage, which was isolated in Wuhan, China, on December 24, 2019 (strain Wuhan-Hu-1, GISAID ID: EPI_ISL_402123) (Wu et al., 2020). Focusing on the evolutionary scenario leading to the emergence of Omicron, the B.1 lineage, which has acquired the D614G mutation in the spike (S) protein (Hou et al., 2020; Korber et al., 2020; Li et al., 2020; Plante et al., 2020; Yurkovetskiy et al., 2020), was first reported on January 24, 2020 (GISAID ID: EPI_ISL_451345). Thereafter, the B.1.1 lineage was first reported in England on February 16, 2020 (GISAID ID: EPI_ISL_466615). The B.1.1 lineage is the common ancestor of Alpha (B.1.1.7 lineage), a prior variant of concern by March 2022, and Omicron (B.1.1.529 lineage), and the Alpha variant caused a large surge of infection worldwide from the fall of 2020 (Davies et al., 2021). Omicron was first reported in South Africa on September 30, 2021 (GISAID ID: EPI_ISL_7971523) (National Institute for Communicable Diseases, 2021).

Soon after the press briefing on Omicron emergence on November 25, 2021 (National Institute for Communicable Diseases, 2021), the virological characteristics of Omicron, currently designated BA.1 (i.e., B.1.1.529.1 lineage, hereafter, the BA.1 lineage is referred to as Omicron in this study), was intensively investigated. For example, Omicron exhibits profound resistance to the humoral immunity induced by vaccination and natural SARS-CoV-2 infection (Cameroni et al., 2021; Cao et al., 2021; Carreño et al., 2021; Cele et al., 2021; Dejnirattisai et al., 2022; Dejnirattisai et al., 2021; Garcia-Beltran et al., 2021; Liu et al., 2021; Meng et al., 2022; Planas et al., 2021; Takashita et al., 2022; VanBlargan et al., 2022). Additionally, we demonstrated that Omicron S is less prone to cleavage by furin, a cellular protease, and exhibits poor fusogenicity (Meng *et al*., 2022; Suzuki et al., 2022). However, it remains unclear why Omicron has spread so rapidly worldwide. In addition, although the explosive infectious spread of Omicron in the human population can be mainly characterized by the virological properties of Omicron S, the mutation(s) in Omicron S that are responsible for its virological characteristics, such as inefficient S cleavage, lower fusogenicity and profound immune resistance, have not been determined.

In this study, we first demonstrate that the representative characteristics of Omicron S, such as immune resistance, poor S cleavage efficiency and poor fusogenicity, are determined by its receptor-binding domain (RBD). By molecular phylogenetic analysis, we show that the acquisition of the S375F mutation in the Omicron RBD is closely associated with the explosive spread of Omicron. Moreover, we experimentally demonstrate that the S375F mutation is critical for the virological properties of Omicron S, namely, the attenuation of S cleavage efficiency and fusogenicity. Furthermore, we elucidate how the attenuated S cleavage and fusogenicity are conferred by the S375F mutation.

## Results

### Omicron RBD determines the virological features of Omicron

To determine the mutation(s) responsible for the virological features of Omicron, we prepared a series of expression plasmids for the Omicron S-based chimeric mutants swapped with the N-terminal domain (NTD) and/or RBD of B.1 (D614G-bearing strain) S (**Figure 1A**). Pseudovirus experiments showed that the pseudovirus with the B.1 RBD-bearing Omicron S [Omicron S/B.1 S_RBD (spike 4 in **Figure 1A**) and Omicron S/B.1 S_NTD+RBD (spike 5)] exhibited increased infectivity compared to that with Omicron S (spike 2) (**Figure 1B**). Western blot analysis showed that the cleavage efficacy of Omicron S was lower than that of B.1 S, which is consistent with our recent studies (Meng *et al*., 2022; Suzuki *et al*., 2022; Yamasoba et al., 2022) (lanes 1 and 2 in **Figures 1C and 1D**). However, the chimeric Omicron S proteins bearing the B.1 RBD (spikes 4 and 5) showed increased cleavage efficacy (**Figures 1C and 1D**). Although the surface expression levels of a series of S chimeras bearing the B.1 domains (spikes 3-5) were lower than those of Omicron S chimeras (**Figure 1E**), a cell-based fusion assay (Motozono et al., 2021; Saito et al., 2022; Suzuki *et al*., 2022; Yamasoba *et al*., 2022) revealed that the fusogenicity of the B.1 RBD-bearing Omicron S was significantly higher than that of the parental Omicron S (**Figure 1F**). To verify the importance of the RBD for the phenotype of Omicron S, we performed reversal experiments based on the B.1 S [B.1 S/Omicron S_RBD (spike 6) in **Figure 1A**]. Corresponding to the results for Omicron S, the pseudovirus infectivity (**Figure 1B**), S cleavage efficacy (**Figures 1C and 1D**), and fusogenicity (**Figures 1F**) of the Omicron RBD-harboring S [B.1 S/Omicron S_RBD (spike 6)] were attenuated compared to those of parental B.1 S. These results suggest that the RBD of Omicron S determines the virological characteristics of Omicron.

**Figure 1.**
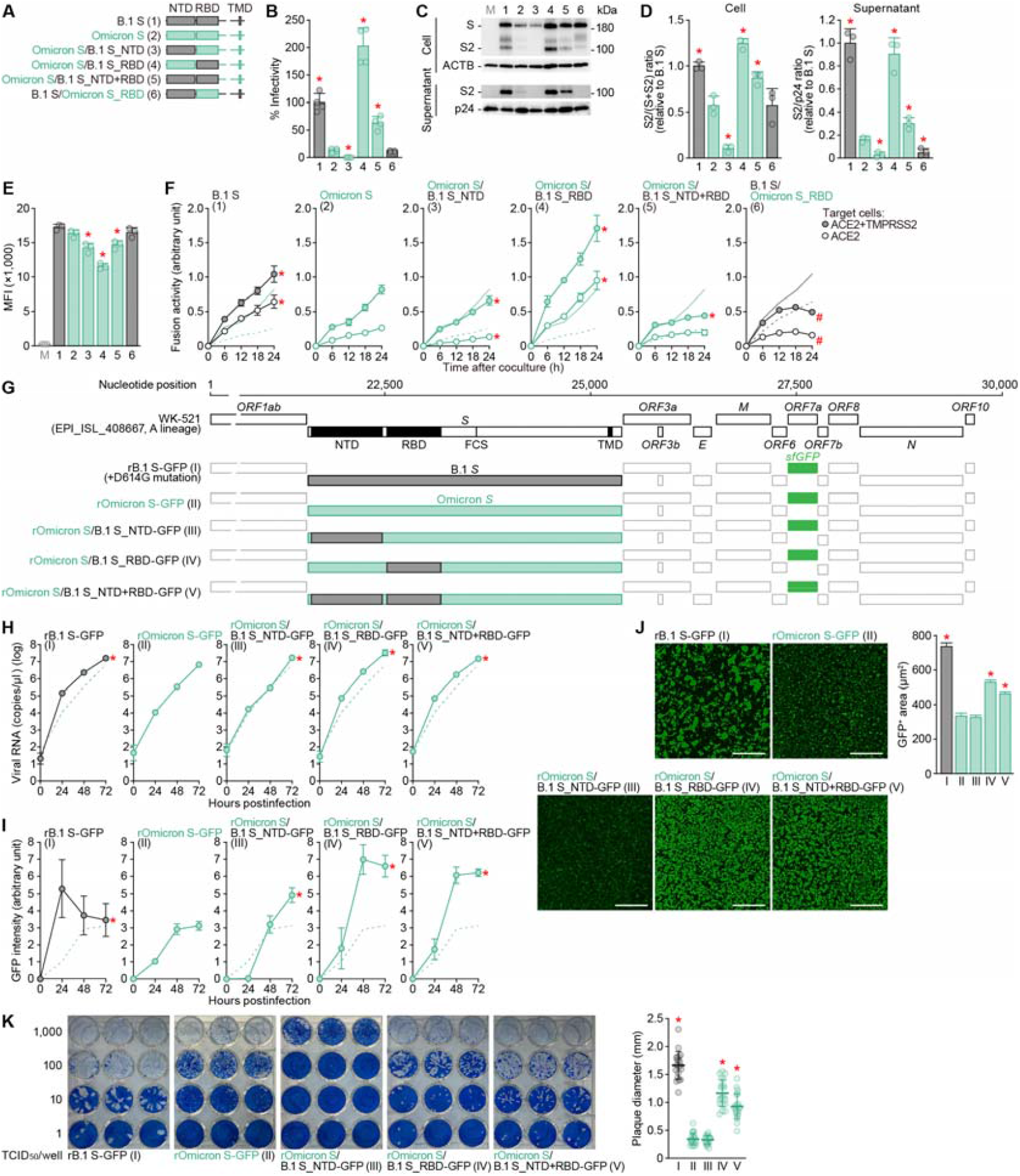
Virological properties conferred by the Omicron RBD. (**A**) Scheme of S chimeras used in this study. The numbers in parentheses are identical to those in **Figures 1B–1E and 2**. NTD, N-terminal domain; RBD, receptor-binding domain; TMD, transmembrane domain. (**B**) Pseudovirus assay. HIV-1-based reporter viruses pseudotyped with SARS-CoV-2 S chimeras (summarized in **Figure 1A**) were prepared. The pseudoviruses were inoculated into HOS-ACE2/TMPRSS2 cells at 1,000 ng HIV-1 p24 antigen, and the percentages of infectivity compared to that of the virus pseudotyped with B.1 S (spike 1) are shown. (**C and D**) Western blot. Representative blots of S-expressing cells and supernatants (**C**) and quantified band intensity (the ratio of S2 to the full-length S plus S2 proteins for “cell”; the ratio of S2 to HIV-1 p24 for “supernatant”) (**D**) are shown. M, mock (empty vector-transfected). (**E**) Flow cytometry. The summarized results of the surface S expression are shown. MFI, mean fluorescent intensity; M, mock (empty vector-transfected). (**F**) SARS-CoV-2 S-based fusion assay. The fusion activity was measured as described in the STAR⍰METHODS, and fusion activity (arbitrary units) is shown. For the target cells, HEK293 cells expressing ACE2 and TMPRSS2 (filled) and HEK293 cells expressing ACE2 (open) were used. The results for B.1 S or Omicron S are shown in other panels as black and green lines, respectively. The results in HEK293-ACE2/TMPRSS2 cells and HEK293-ACE2 cells are shown as normal or broken lines, respectively. (**G**) Scheme of the S-chimeric recombinant SARS-CoV-2 used in this study. FCS, furin cleavage site. The backbone is SARS-CoV-2 strain WK-521 (GISAID ID: EPI_ISL_408667, A lineage) (Torii *et al*., 2021). Note that the *ORF7a* gene is swapped with the *sfGFP* gene. The numbers in parentheses are identical to those in **Figures 1H–1K**. (**H-J**) SARS-CoV-2 infection. VeroE6/TMPRSS2 cells were infected with a series of chimeric recombinant SARS-CoV-2 (shown in **G**) at multiplicity of infection (m.o.i.) 0.01. Viral RNA in the supernatant (**H**) and GFP intensity (**I**) were measured using routine techniques. The result for Omicron (virus II) is shown in other panels as a broken green line. (**J**) Syncytium formation. Left, GFP-positive area at 48 h.p.i. Scale bar, 500 μm. Right, summarized results. I, n=6,483 cells; II, n=5,393 cells; III, n=8,704 cells; IV, n=13,188 cells; and V, n=12,749 cells. Representative images are shown in **Figure S1**. (**K**) Plaque assay. Left, representative figures. Right, summary of the plaque diameters (20 plaques per virus). Data are expressed as the mean with SD (**B, D-F, H and K**) or the median with 95% confidence interval (CI) (**J**). Assays were performed in quadruplicate (**B, H**) or triplicate (**D-F**). Each dot indicates the result of an individual replicate (**B, D and E**) or an individual plaque (**K**). Statistically significant differences (*P<0.05) versus Omicron S (pseudovirus 2 for **B, D and E**, virus II for **J and K**) were determined by two-sided Student’s t test (**B and E**), two-sided paired t test (**D**), or two-sided Mann–Whitney U test (**J and K**). In **F, H and I**, statistically significant differences [*familywise error rates (FWERs)<0.05] versus Omicron (spike 2 or virus II) (except for the rightmost panel in **F**) or B.1 (spike 1 or virus I) (rightmost panel in **F**) through timepoints were determined by multiple regression. FWERs were calculated using the Holm method. See also **Figure S1**.

To further investigate the impact of the Omicron S RBD, we generated a series of recombinant chimeric SARS-CoV-2 strains by reverse genetics (**Figure 1G**) (Torii et al., 2021). As shown in **Figure 1H**, the growth of rOmicron S-GFP (virus II) and rOmicron S/B.1 S_NTD (virus III) was lower than that of rB.1 S-GFP (virus I). However, the recombinant viruses bearing the B.1 RBD [rOmicron S/B.1_RBD-GFP (virus IV) and rOmicron S/B.1 S_NTD+RBD-GFP (virus V)] replicated more efficiently than rOmicron S-GFP (virus II) in VeroE6/TMPRSS2 cells (**Figure 1H**). Additionally, we measured the GFP intensity in infected cell cultures using routine procedures and showed that the GFP intensity of the cells infected with the recombinant viruses bearing the B.1 RBD was significantly higher than that of the cells infected with rOmicron S-GFP (virus II) (**Figures 1I and S1**). These data suggest that the RBD of Omicron S attenuates viral growth capacity in cell cultures. To evaluate the fusogenicity of the chimeric viruses, we measured the level of GFP-positive cells. As shown in **Figure 1J**, the GFP-positive area of the cells infected with the recombinant viruses at 48 hours post infection (h.p.i.) was significantly larger for viruses bearing the B.1 RBD [rOmicron S/B.1_RBD-GFP (virus IV) and rOmicron S/B.1 S_NTD+RBD-GFP (virus V)] than for rOmicron-GFP (virus II). Consistent with the results using S-expressing cells (**Figure 1F**), these findings suggest that the Omicron RBD attenuates viral fusogenicity. Moreover, the plaques formed by infection with rOmicron S/B.1 S_RBD-GFP (virus IV) and rOmicron S/B.1 S_NTD+RBD-GFP (virus V) were significantly larger than those formed by rOmicron S-GFP virus (virus II), while the plaques formed by rOmicron S-GFP (virus II) and rOmicron S/B.1 S_NTD-GFP (virus III) were comparable (**Figure 1K**). Altogether, these results suggest that the RBD of Omicron S determines the virological features of this viral lineage, such as the attenuation of pseudovirus infectivity, S1/S2 cleavage efficacy and fusogenicity, of Omicron.

### Omicron RBD mainly determines the immune resistance of Omicron

We next assessed the domains of Omicron S that are associated with the profound immune resistance of Omicron (Cameroni *et al*., 2021; Cao *et al*., 2021; Carreño *et al*., 2021; Cele *et al*., 2021; Dejnirattisai *et al*., 2022; Dejnirattisai *et al*., 2021; Garcia-Beltran *et al*., 2021; Liu *et al*., 2021; Meng *et al*., 2022; Planas *et al*., 2021; Takashita *et al*., 2022; VanBlargan *et al*., 2022). Because the swapping of Omicron S with B.1 S NTD (Omicron S/B.1 S_NTD, spike 3) severely decreased pseudovirus infectivity (**Figure 1B**), we performed neutralization assays using pseudoviruses with Omicron RBD-bearing B.1 S [Omicron S/B.1 S_RBD (spike 4) and Omicron S/B.1 S_NTD+RBD (spike 5) as well as the S proteins of Omicron (spike 2), Delta and B.1 (spike 1) (the list of sera used is shown in **Table S1**). Consistent with recent studies (Cameroni *et al*., 2021; Cao *et al*., 2021; Carreño *et al*., 2021; Cele *et al*., 2021; Dejnirattisai *et al*., 2022; Dejnirattisai *et al*., 2021; Garcia-Beltran *et al*., 2021; Liu *et al*., 2021; Meng *et al*., 2022; Planas *et al*., 2021; Takashita *et al*., 2022; VanBlargan *et al*., 2022), Omicron S (spike 2) was highly resistant to the vaccine sera [BNT162b2 (**Figure 2A**) and mRNA-1273 (**Figure 2B**)] as well as convalescent sera from individuals infected with early pandemic virus (collected before May 2020) (**Figure 2C**) or with the Delta variant (**Figure 2D**). In the cases of pseudoviruses with Omicron S/B.1 S_RBD (spike 4) and Omicron S/B.1 S_NTD+RBD (spike 5), these pseudoviruses were significantly more sensitive to vaccine sera (**Figures 2A and 2B**) and convalescent sera obtained from early pandemic virus-infected patients than Omicron S (spike 2) (**Figure 2C**). These results suggest that the RBD of Omicron S is closely associated with its pronounced resistance to antiviral humoral immunity elicited by vaccination or previous SARS-CoV-2 infection. Moreover, we used convalescent sera from hamsters infected with B.1.1 (note that the S gene sequences of B.1 and B.1.1 are identical) and Omicron for the assay. As shown in **Figure 2E**, Omicron S (spike 2) was completely resistant to the B.1.1 convalescent sera, while it was sensitive to the Omicron convalescent sera. Notably, the chimeric Omicron S bearing the B.1 RBD [Omicron S/B.1 S_RBD (spike 4) and Omicron S/B.1 S_NTD+RBD (spike 5)] exhibited the opposite results: these chimeric pseudoviruses were sensitive to the B.1.1 convalescent sera (**Figure 2E**) but completely resistant to the Omicron convalescent sera (**Figure 2F**). These results further suggest that the Omicron RBD determines its immune resistance and can be a remarkable antigen for humoral immunity. However, we found that Omicron S/B.1 S_NTD+RBD (spike 5) is significantly more sensitive to antisera than Omicron S/B.1 S_RBD (spike 4) (**Figures 2A–2C and 2E**). These findings suggest that mutations in the NTD of Omicron S are also partly associated with the immune resistance of Omicron S.

**Figure 2.**
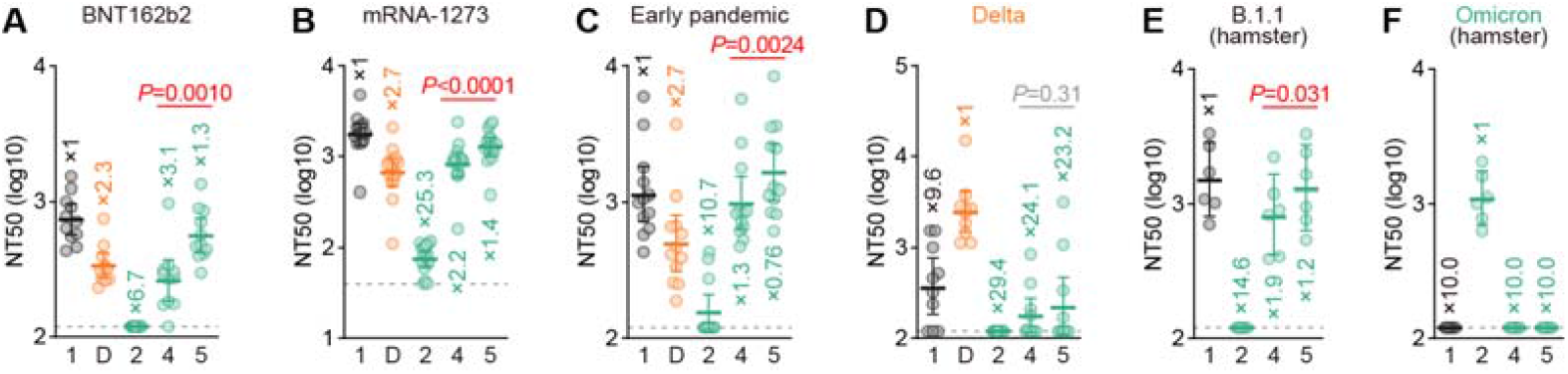
Immune resistance conferred by the Omicron RBD. Neutralization assays were performed with pseudoviruses harboring a series of S protein sequences (summarized in **Figure 1A**). The numbers are identical to those in **Figure 1A**. D, Delta variant. Vaccinated sera [BNT162b2 (**A**, 11 donors); or mRNA-1273 (**B**, 16 donors)], convalescent sera of individuals infected with an early pandemic virus (before May 2020) (**C**, 12 donors), or Delta (**D**, 10 donors) and convalescent sera of hamsters infected with B.1.1 (**E**, 6 hamsters) or Omicron (**F**, 6 hamsters) were used. The list of sera used in this experiment is shown in **Table S1**. Each serum sample was analyzed in triplicate to determine the 50% neutralization titer (NT50). Each dot represents one NT50 value, and the geometric mean and 95% CI are shown. The numbers indicate the fold changes of resistance versus each antigenic variant. Horizontal gray lines indicate the detection limit of each assay (120 for **A and C-F**; 40 for **B**). Statistically significant differences between spikes 4 and 5 were determined by a two-sided Wilcoxon signed-rank test. See also Table S1.

### S S375F mutation increases binding affinity to human ACE2

In the RBD (residues 319-541), 12 substitutions were uniquely found in Omicron S, while the other 3 substitutions (K417N, T478K and N501Y) were commonly detected in the other variants (**Figure 3A**) (Meng *et al*., 2022). To determine the residue(s) responsible for the virological phenotype of Omicron, we prepared a series of B.1 S RBD point mutants that bear the respective mutations of Omicron and conducted screening experiments based on a yeast surface display assay (Dejnirattisai *et al*., 2022; Kimura et al., 2022; Motozono *et al*., 2021; Yamasoba *et al*., 2022; Zahradnik et al., 2021). As shown in **Figure 3B** (left panel), compared to the RBD of parental (i.e., B lineage-based) S, the K_D_ values of the G339D, N440K and S477N mutants significantly decreased, while those of the S375F, S371L/S373P/S375F, G496S and Y505H mutants significantly increased. Since the ACE2 binding affinity of Omicron S is lower than that of the RBD of ancestral (including B.1 lineage) SARS-CoV-2 (Dejnirattisai *et al*., 2022; Yamasoba *et al*., 2022), these data suggest that the S375F, G496S and Y505H substitutions are closely associated with the phenotypes of Omicron.

**Figure 3.**
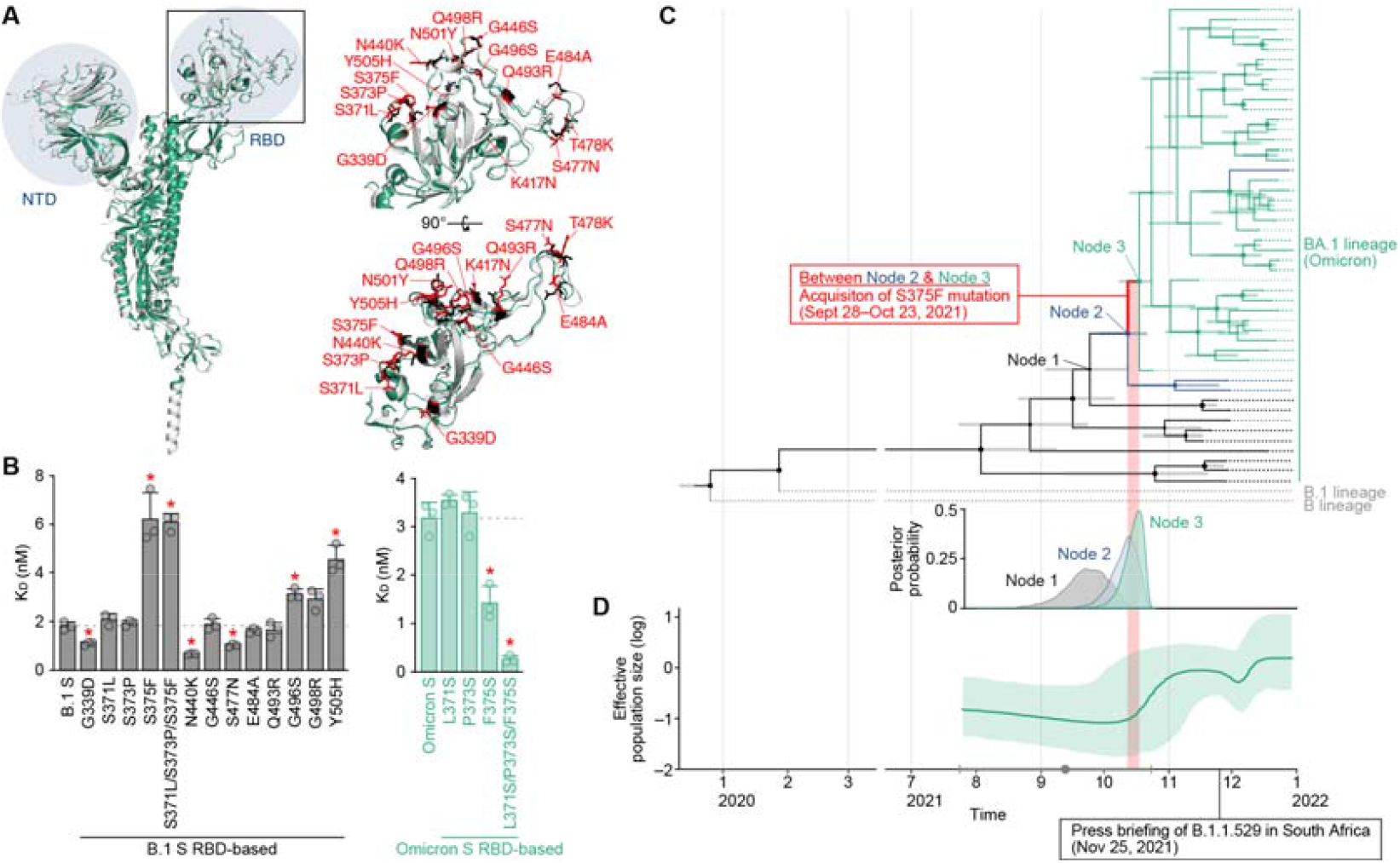
Mutations in the Omicron RBD and the evolution of Omicron. (**A**) Structural insights into the mutations in the Omicron RBD. Left, overlaid crystal structures of SARS-CoV-2 B.1 S (PDB: 7KRQ) (Zhang et al., 2021) (white) and Omicron S (PDB: 7T9J) (Mannar et al., 2022) (green) are shown. The NTD and RBD are indicated in blue. The region in the RBD indicated by a square is enlarged in the top right panel. Right, mutated residues in the RBD. The residues in B.1 S and Omicron S are shown in black and red, and the mutations in Omicron S are indicated. (**B**) ACE2 binding affinity of a series of SARS-CoV-2 S RBD (residues 336-528) mutants tested by yeast surface display. The K_D_ values of the binding of the SARS-CoV-2 S RBD expressed on yeast to soluble ACE2 are shown. (**C and D**) Evolution of Omicron. (**C**) Top, a time tree of 48 Omicron variants and two outgroups (B and B.1 lineages). The same tree annotated with the GISAID ID, PANGO lineage and sampling date at each terminal node is shown in **Figure S2**. Green, Omicron variants containing the S371L, S373P and S375F mutations; blue, Omicron variants containing the S371L and S373P mutations; black, Omicron variants without the S371L/S373P/S375F mutations; and gray, the two outgroups (B and B.1 lineages). The bars on each internal node indicate the 95% highest posterior density (HPD) interval of the estimated time. Note that “Node 1” corresponds to the time to before the emergence of the S371L and S373P mutations; “Node 2” corresponds to the time after the acquisition of the S371L and S373P mutations and before the emergence of the S375F mutations; and “Node 3” corresponds to the fixation time of the S371L/S373P/S375F mutations in the Omicron variants. The estimated time of each node is as follows: Node 1, September 23, 2021 (95% HPD September 2, 2021 to October 13, 2021); Node 2, October 11, 2021 (95% HPD September 28, 2021 to October 21, 2021); and Node 3, October 16, 2021 (95% HPD October 7, 2021 to October 23, 2021). Bottom, distribution of the posterior probability of the time to the tMRCA of Node 1 (black), Node 2 (blue), and Node 3 (green). (**D**) Bayesian skyline plot showing the history of the effective population size of 48 Omicron variants. The 95% HPD is shaded in red. The dot (in gray) indicates the estimated tMRCA of the 48 variants (September 12, 2021), and the error bar (in gray) indicates the lower (July 24, 2021) and upper (October 23, 2021) boundaries of the 95% HPD tMRCA. In **B**, the data are expressed as the mean with SD. The assay was performed in triplicate, and each dot indicates the result of an individual replicate. The horizontal broken lines indicate the value of B.1 S (left) and Omicron S (right), respectively. Statistically significant differences (*P<0.05) versus B.1 S (left) or Omicron S (right) were determined by two-sided Student’s t tests, and FWERs were calculated using the Holm method. In **C** and **D**, the timeline on the x-axis was shared, and the time of S375F emergence (i.e., between “Node 2” and “Node 3” in **C**) is shaded in red. **See also Figure S2**.

### Evolution of Omicron is closely associated with the acquisition of the S S375F mutation

The S375F, G496S and Y505H mutations in the S protein were almost exclusively detected in the Omicron variants (**Table S2**). To infer the evolutionary sequence of the emergence of these mutations in the lineage of Omicron, we generated a time tree of 48 Omicron genomes detected in 2021 (for more detail, see STAR⍰METHODS) (**Figures 3C and S2**). The G496S and Y505H mutations were detected in all sequences used in this analysis, suggesting that these two mutations were acquired in the common ancestor of all Omicron variants reported so far. In contrast, the S371L, S373P and S375F mutations were not included in the older Omicron sequences (shown in black in **Figures 3C and S2**). Although the emergence times of S371L and S373P cannot be estimated independently, our analysis assumed that the S371L and S373P mutations were first acquired between Node 1 [95% highest posterior density (HPD): September 2, 2021 to October 13, 2021] and Node 2 (95% HPD: September 28, 2021 to October 21, 2021) in **Figure 3C**, based on the estimated time to the most recent common ancestor (tMRCA). The S375F mutation emerged thereafter, between Node 2 and Node 3 (95% HPD: October 7, 2021 to October 23, 2021) (**Figure 3C**). Interestingly, the Bayesian skyline plot of the 48 Omicron genomes suggested that the effective population size of Omicron increased just after the acquisition of the S375F substitution (**Figure 3D**). These data suggest that the emergence of the S375F mutation was a crucial event to trigger the massive spread of the Omicron variants in the human population.

To verify the possibility that the S375F mutation is crucial for the phenotype of Omicron, we performed yeast binding assays using the RBD of Omicron S. As shown in **Figure 3B** (right panel), the F375S and L371S/P373S/F375S mutations in the RBD of Omicron S significantly increased the binding affinity to human ACE2. Overall, these observations suggest that the three substitutions at positions 371, 373 and 375, and particularly the S375F substitution, determine the virological phenotype of Omicron.

### S S375F mutation determines the virological features of Omicron

To investigate the impact of the S375F mutation, we prepared pseudoviruses with a series of Omicron S-based mutants (**Figure 4A**). Corresponding to the yeast surface display assay (**Figure 3B**), the gain-of-function assay showed that pseudovirus infectivity was clearly increased by the insertion of the F375S mutation (spikes 9 and 11-13 in **Figure 4A**) in Omicron S (**Figure 4B, top**). Western blot analysis showed that the S1/S2 cleavage efficacy was also rescued by the F375S mutation (**Figures 4C and 4D, top**). Although the surface S expression level was decreased by this mutation (**Figure 4E, top**), a cell-based fusion assay demonstrated that the F375S mutation significantly increased the efficacy of SARS-CoV-2 S-mediated cell–cell fusion (**Figure 4F, top**). Conversely, a loss-of-function assay based on the B.1 S showed that the S375F mutation (spikes 16 and 18-20) decreased pseudovirus infectivity (**Figure 4B, bottom**), S cleavage efficacy (**Figures 4C and 4D, bottom**) and fusion activity (**Figure 4F, bottom**). These results suggest that the S375F mutation in Omicron S is responsible for its virological phenotypes. However, the S371L/S373P/S375F mutations did not affect the sensitivity to the antiviral humoral immunity elicited by vaccination and infection (**Figure S3**), suggesting that these mutations are not associated with the immune resistant phenotype of Omicron.

**Figure 4.**
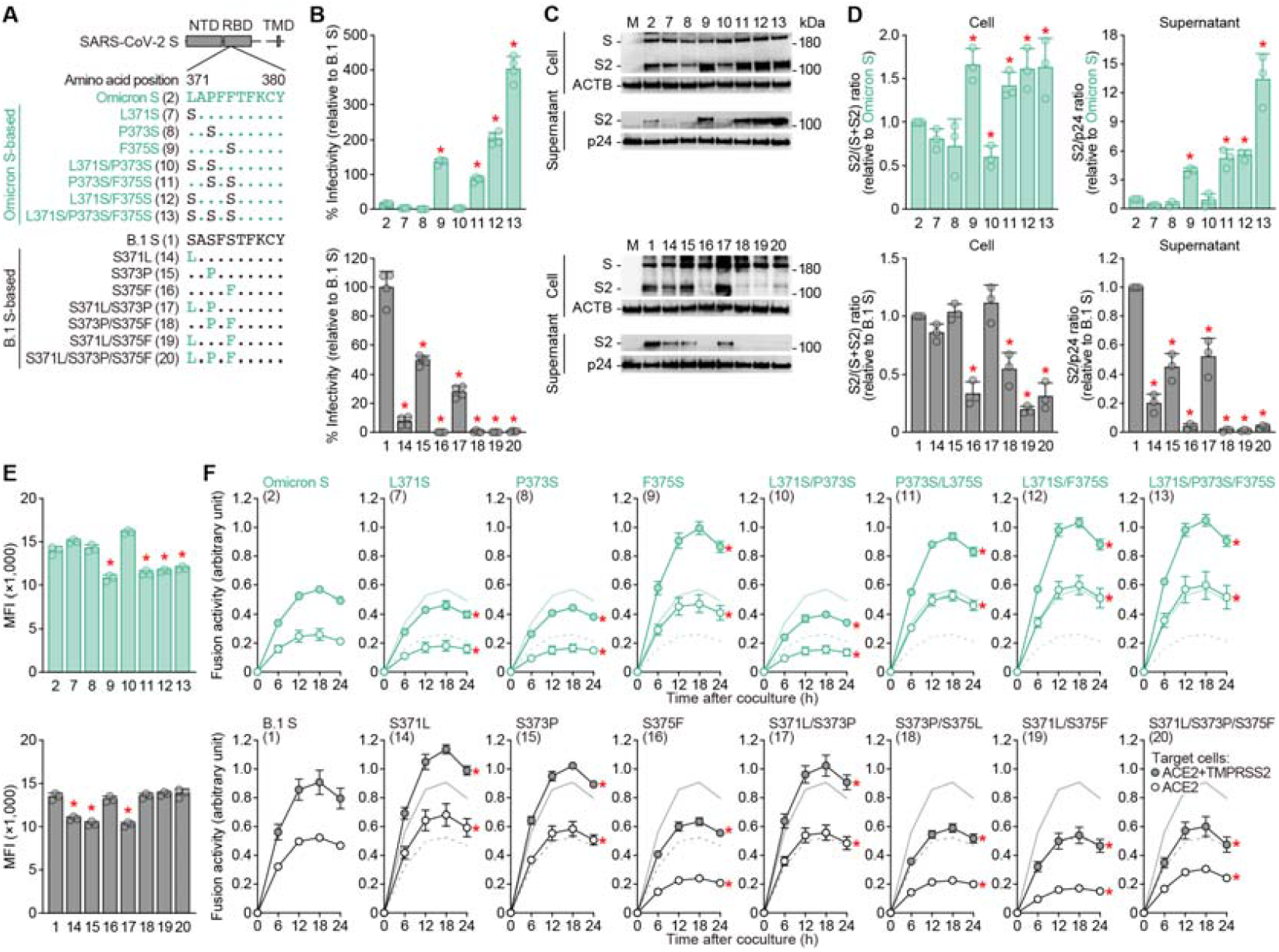
Virological features conferred by the S S375F mutation. (**A**) Scheme of the S mutants used in this study. The numbers in parentheses are identical to those in **Figures 4B–4F and S3**. (**B**) Pseudovirus assay. HIV-1-based reporter viruses pseudotyped with SARS-CoV-2 S mutants (summarized in **A**) were prepared. The pseudoviruses were inoculated into HOS-ACE2/TMPRSS2 cells at 1,000 ng HIV-1 p24 antigen, and the percent infectivity compared to that of the virus pseudotyped with Omicron S (spike 2, top) or B.1 S (spike 1, bottom) are shown. (**C and D**) Western blot. Representative blots of S-expressing cells and supernatants (**C**) and quantified band intensity (the ratio of S2 to the full-length S plus S2 proteins for “cell”; the ratio of S2 to HIV-1 p24 for “supernatant”) (**D**) are shown. M, mock (empty vector-transfected). (**E**) Flow cytometry. The summarized results of the surface S expression are shown. (**F**) SARS-CoV-2 S-based fusion assay. The fusion activity was measured as described in STAR⍰METHODS, and fusion activity (arbitrary units) is shown. For the target cells, HEK293 cells expressing ACE2 and TMPRSS2 (filled) and HEK293 cells expressing ACE2 (open) were used. The results for Omicron S (top) or B.1 S (bottom) are shown in other panels as green and black lines, respectively. The results in HEK293-ACE2/TMPRSS2 cells and HEK293-ACE2 cells are shown as normal and broken lines, respectively. Data are expressed as the mean with SD. Assays were performed in quadruplicate (**B**) or triplicate (**D-F**). In **B, D and E**, each dot indicates the result of an individual replicate. Statistically significant differences (*P<0.05) versus the respective parental S [Omicron S (pseudovirus 2, top panels) or B.1 S (spike 1, bottom panels)] were determined by two-sided Student’s t test (**B and E**) or two-sided paired t test (**D**). In **F**, statistically significant differences (*FWERs<0.05) versus the respective parental S [Omicron S (spike 2, top panels) or B.1 S (spike 1, bottom panels)] through timepoints were determined by multiple regression. FWERs were calculated using the Holm method. **See also** Figure S3.

To further assess the impact of the S375F mutation, we generated two additional recombinant chimeric SARS-CoV-2 strains, B.1 S S375F-GFP (virus VI) and Omicron S F375S-GFP (virus VII) (**Figure 5A**). Although the mutation at position 375 did not affect the viral RNA load in the culture supernatant of infected VeroE6/TMPRSS2 cells (**Figure 5B**), the GFP intensity in infected VeroE6/TMPRSS2 cells was modulated by the mutation at the position 375 of the S protein: the S375F mutation in the B.1 S backbone decreased the GFP intensity, while the F375S mutation in the Omicron S backbone increased the intensity (**Figures 5C and S1**). Additionally, quantitative fluorescence microscopy showed that the GFP-positive area of B.1 S S375F-GFP (virus VI) was significantly lower than that of parental B.1 S-GFP (virus I), while that of Omicron S F375S-GFP (virus VII) was significantly higher than that of parental Omicron S-GFP (virus II) (**Figure 5D**). Moreover, plaque assays showed that the plaques formed by infection with B.1 S S375F-GFP (virus VI) were significantly smaller than those formed by B.1 S-GFP (virus I), while conversely, the plaque size was increased by the insertion of the F375S mutation in Omicron S (**Figure 5E**). Altogether, these results suggest that the S375F mutation in the Omicron S protein determines the virological characteristics (decreased infectivity, decreased S1/S2 cleavage efficacy, and decreased fusogenicity) of Omicron.

**Figure 5.**
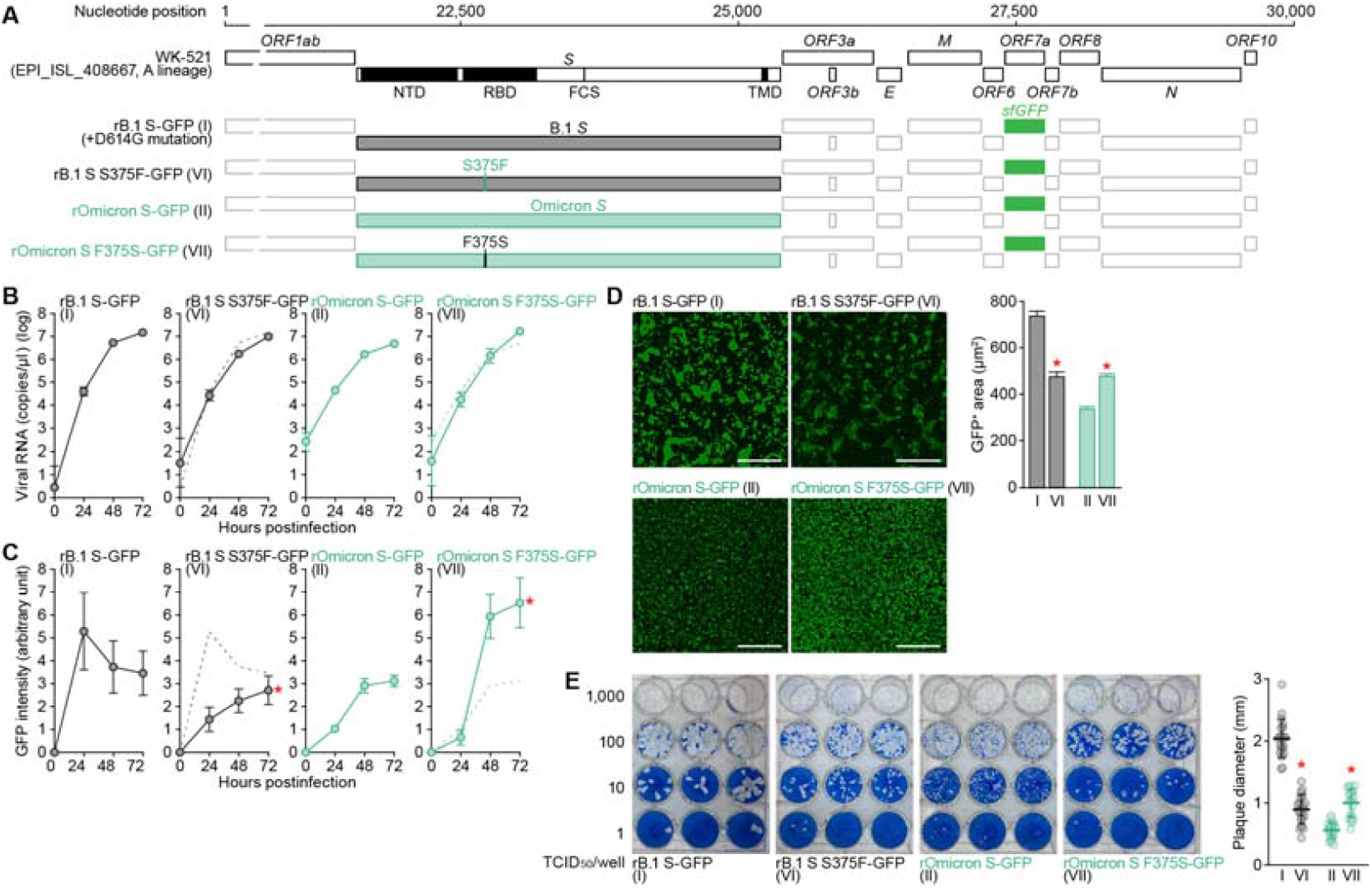
Effect of the S S375F mutation on viral growth dynamics. (**A**) Scheme of the S-chimeric recombinant SARS-CoV-2 used in this study. The numbers in parentheses are identical to those in **Figures 5B–5E**. (**B-D**) SARS-CoV-2 infection. VeroE6/TMPRSS2 cells were infected with a series of S-chimeric recombinant SARS-CoV-2 (summarized in **A**) at an m.o.i. 0.01. The viral RNA in the supernatant (**B**) and GFP intensity (**C**) were measured routinely. The results for the respective parental S are shown in other panels as broken green lines. Assays were performed in quadruplicate (**B and C**). (**D**) Syncytium formation. Left, GFP-positive area at 48 h.p.i. Scale bar, 500 μm. Right, summarized results. I, n=6,483 cells; VI, n=2,780 cells; II, n=5,393 cells; and VII, 12,857 cells. The results for B.1-GFP (virus I) and Omicron-GFP (virus II) in **C** and **D** (right) are identical to those shown in **Figures 1I and 1J** (right). Representative images are shown in **Figure S1**. (**E**) Plaque assay. Left, representative figures. Right, summary of the plaque diameters (20 plaques per virus). Each dot indicates the result of an individual plaque. Data are expressed as the mean with SD (**B and E**) or the median with 95% CI (**D**). In **B and C**, statistically significant differences (*FWERs<0.05) versus Omicron-GFP (virus II) through timepoints were determined by multiple regression. FWERs were calculated using the Holm method. In **D and E**, statistically significant differences (*P<0.05) versus Omicron-GFP (virus II) were determined by a two-sided Mann–Whitney U test. See also **Figure S1.**

### The F375-H505 pi-pi interaction contributes to the virological phenotype of Omicron

Here, we experimentally demonstrated that the S375F mutation determines the virological properties of Omicron (**Figures 4 and 5**). Additionally, molecular phylogenetic analysis suggested that the emergence of this mutation was closely associated with the explosive growth of Omicron in the human population (**Figures 3C and 3D**). However, it remains unclear how the S375F mutation contributes to the phenotype of Omicron at the molecular level. We addressed this question using a structural biology approach. As shown in **Figure 6A** (top), we predicted that the F375 residue in a fully closed Omicron S trimer can form a pi-pi interaction, a sort of dispersion by van der Waals forces between aromatic residues [reviewed in (Martineza and Iverson, 2012)], with the H505 residue in another S protein in the same trimer. Because residue 375 in the B.1.1 S protein is a serine, the pi-pi interaction cannot be formed (**Figure 6A, bottom**). To address the hypothesis that the F375-H505-mediated inter-protomer pi-pi interaction contributes to the phenotype of Omicron, we prepared the Omicron S H505A mutant in which an aromatic side chain at position 505 is disrupted. Western blot analysis showed that the S cleavage efficacy of Omicron S was increased by the insertion of the H505A mutation (**Figure 6B**). To further test this possibility, the residues at position 375 of B.1 S were substituted with amino acids bearing aromatic side chains (i.e., F, Y and H). Similar to the S375F mutant, the B.1 S mutants bearing the S375Y or S375H mutation showed decreased cleavage efficacy of the S protein (**Figure 6C**). These results further suggest that the inter-protomer pi-pi interaction is formed between Y505 and S375F/Y/H. Moreover, the insertion of the Y505A mutation in the B.1 S bearing the S375F/Y/H mutation (i.e., the disruption of the aromatic residue at position 505) rescued the S cleavage efficacy (**Figure 6C**).

**Figure 6.**
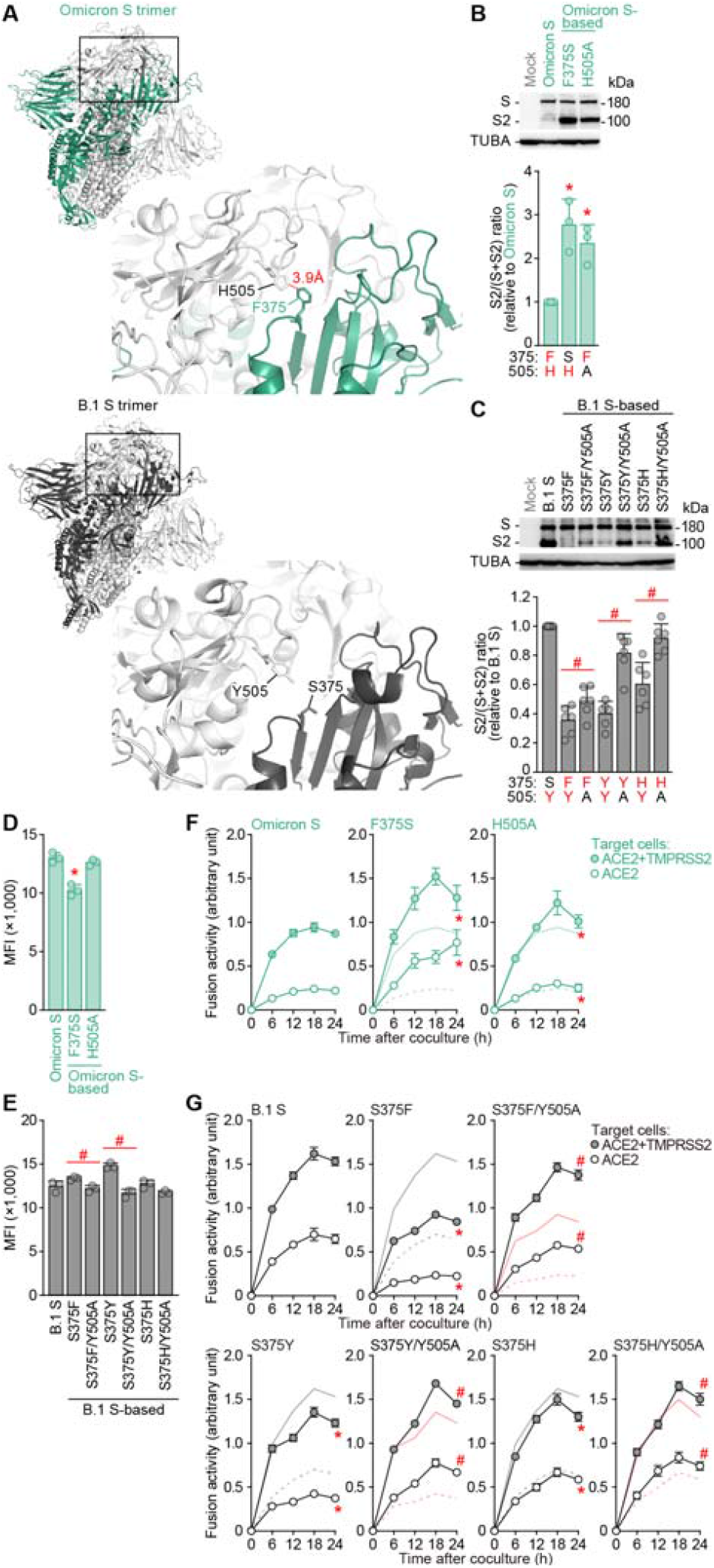
Effect of the pi-pi interaction between 375F and 505H. (**A**) Structural insights into the SARS-CoV-2 S trimer. Top, the structure of the Omicron S trimer (PDB: 7T9J) (Mannar *et al*., 2022) reconstructed as described in the STAR⍰METHODS. Bottom, crystal structure of the B.1 S trimer (PDB: 7KRQ) (Zhang *et al*., 2021). The regions indicated in squared are enlarged in the bottom right panels. In the enlarged panels, the residues at position 375 [F in an Omicron S monomer indicated in green (top); S in a B.1 S monomer indicated in black (bottom)] and 505 [H in an Omicron S monomer indicated in white (top); Y in a B.1 S monomer indicated in white (bottom)] are shown. The putative pi-pi interaction between F375 and H505 in the Omicron S trimer is indicated in red (3.9 Å). (**B and C**) Western blot. Representative blots of S-expressing cells (top) and quantified band intensity (the ratio of S2 to the full-length S plus S2 proteins) (bottom) are shown. In the bottom panels, the residues at positions 375 and 505 are indicated, and aromatic residues (F, H or Y) are indicated in red. (**D and E**) Flow cytometry. The summarized results of the surface S expression are shown. (**F and G**) SARS-CoV-2 S-based fusion assay. The fusion activity was measured as described in the STAR⍰METHODS, and fusion activity (arbitrary units) is shown. For the target cells, HEK293 cells expressing ACE2 and TMPRSS2 (filled) and HEK293 cells expressing ACE2 (open) were used. In **F**, normal lines, Omicron S with HEK293-ACE2/TMPRSS2 cells; broken lines, Omicron S with HEK293-ACE2 cells. In the panels of S375F, S375Y and S375H in **G**, normal black lines, B.1 S using HEK293-ACE2/TMPRSS2 cells; broken black lines, B.1 S using HEK293-ACE2 cells; normal red lines. In the panels for S375F/Y505A, S375Y/Y505A and S375H/Y505A in **G**, normal red line, the result for the respective mutant without the Y505A mutation using HEK293-ACE2/TMPRSS2 cells; broken red line, the result for the respective mutant without the Y505A mutation using HEK293-ACE2 cells. Data are expressed as the mean with SD. Assays were performed in triplicate (**B, D-G**) or sextuplicate (**C**). In **B-E**, each dot indicates the result of an individual replicate. Statistically significant differences versus Omicron S (*P<0.05) and between the mutant with and without the Y505A mutation (#P<0.05) were determined by two-sided paired t test (**B and C**) or two-sided Student’s t test (**D and E**). In **F and G**, statistically significant differences versus Omicron S (*FWERs<0.05) or the mutant without the Y505A mutation (#FWERs<0.05) through timepoints were determined by multiple regression. FWERs were calculated using the Holm method.

Finally, we verified the impact of the inter-protein pi-pi interaction on S-mediated fusogenicity. Although the Omicron S F375S mutant exhibited a decreased surface expression level, the H505A mutation did not (**Figure 6D**). In the case of the B.1 S-based mutants, the Y505A mutation decreased the surface expression levels when inserted together with the S375F/Y mutations (**Figure 6E**). Corresponding to the western blot results (**Figure 6B**), the disruption of the pi-pi interaction by F375S and H505A in Omicron S significantly increased fusion activity (**Figure 6F**). Moreover, in the case of the B.1 S-based mutant, the substitution of residue 375 with an aromatic residue (F, Y or H) significantly decreased the fusion activity (**Figure 6G**). However, when we inserted the Y505A substitution in the S375F/Y/H mutants to disrupt the aromatic residue at position 505, fusion activity was significantly increased (**Figure 6G**). Altogether, our results suggest that the inter-protomer pi-pi interaction mediated by the aromatic residues at positions 375 and 505 of the S protein contributes to decreased S cleavage efficacy and fusogenicity.

## Discussion

In the present study, we performed multiscale investigations to unveil the virological characteristics of the Omicron variant of SARS-CoV-2. By using pseudoviruses, a yeast surface display system and the chimeric recombinant SARS-CoV-2 generated by reverse genetics, we showed that the RBD of Omicron S is responsible for the representative virological features of this variant. In particular, the S375F mutation in the RBD of Omicron S determines the characteristic virological properties of Omicron: decreased affinity to ACE2, decreased infectivity, decreased growth efficacy, attenuated efficacy of S cleavage, and reduced fusogenicity. Molecular phylogenetic analysis provided evidence suggesting that the acquisition of the S375F mutation was closely related to the onset of the explosive spread of Omicron in the human population. Furthermore, experiments based on structural biology revealed that the pi-pi interaction mediated by residues of F375 and H505 is responsible for the characteristics of Omicron S.

We provided evidence suggesting that the S375F mutation determines the virological features of Omicron. We also revealed that the nascent pi-pi interaction in the S trimer is established by the F375 and H505 residues and characterizes the Omicron S. Because the Y505H mutation was already acquired in the ancestral Omicron sequences, our results suggest that the acquisition of the S375F mutation during the evolution of Omicron rendered the properties of SARS-CoV-2 S protein to attenuate the S cleavage efficacy and fusogenicity, which led to the explosive spread of Omicron in the human population. The S375F mutation is highly conserved in the Omicron lineage and has not been detected in the other SARS-CoV-2 variants that have emerged to date. However, our data suggest that the substitution of residues possessing an aromatic ring, such as phenylalanine, tyrosine and histidine, at residue 375 can confer Omicron-like properties. Therefore, the emergence of SARS-CoV-2 variants bearing such substitutions at residue 375 should be considered a potential risk for global health.

### Limitations of the study

Here, we showed the importance of the S375F mutation to the virological properties of Omicron; however, the following issues remain unclear. First, although we showed that the S375F mutation determines the virological features of Omicron S, it remains unclear which mutations in Omicron S determine the pronounced immune resistance of Omicron S. Second, in addition to the Omicron BA.1 variant that we focused on this study, another recently emerged Omicron lineage, BA.2, also bears the S375F mutation. However, we have recently shown that the fusogenicity of BA.2 S is significantly higher than that of BA.1 S (Yamasoba *et al*., 2022). Together with the results shown in this study, these observations suggest that BA.2 S acquired certain compensatory mutation(s) that increased fusion efficacy. Further investigations will be needed to unveil the full evolutionary history of the Omicron lineage. Furthermore, the question of why the acquisition of the S375F mutation caused explosive spread despite reducing infectivity in tissue culture, S cleavage efficacy and fusogenicity also needs to be elucidated in detail by further studies.

## Author Contributions

Izumi Kimura, Daichi Yamasoba, Hesham Nasser, Yusuke Kosugi, Kayoko Nagata, Keiya Uriu, Yuri L Tanaka, Ryo Shimizu, Toong Seng Tan, Erika P Butlertanaka, Mako Toyoda, Takashi Irie, Akatsuki Saito and Terumasa Ikeda performed cell culture experiments.

Jiri Zahradnik and Gideon Schreiber performed a yeast surface display assay. Jumpei Ito, Hiroyuki Asakura, Kenji Sadamasu and Kazuhisa Yoshimura performed viral genome sequencing analysis.

Takamasa Ueno, Akifumi Takaori-Kondo and Kotaro Shirakawa contributed clinical sample collection.

Jumpei Ito performed statistical analyses.

Jiri Zahradnik and Yusuke Kosugi performed structural analyses.

Jiaqi Wu and So Nakagawa performed molecular phylogenetic analyses.

Mako Toyoda, Kotaro Shirakawa, Takashi Irie, Akatsuki Saito, So Nakagawa, Terumasa Ikeda and Kei Sato designed the experiments and interpreted the results.

Kei Sato wrote the original manuscript.

All authors reviewed and proofread the manuscript.

The Genotype to Phenotype Japan (G2P-Japan) Consortium contributed to the project administration.

## Acknowledgments

We would like to thank all members belonging to The Genotype to Phenotype Japan (G2P-Japan) Consortium. We thank Dr. Kenzo Tokunaga (National Institute for Infectious Diseases, Japan) and Dr. Jin Gohda (The University of Tokyo, Japan) for providing reagents. The super-computing resource was provided by Human Genome Center at The University of Tokyo.

This study was supported in part by AMED Research Program on Emerging and Re-emerging Infectious Diseases (20fk0108268, to Akifumi Takaori-Kondo; 20fk0108517, to Akifumi Takaori-Kondo; 20fk0108146, to Kei Sato; 20fk0108270, to Kei Sato; 20fk0108413, to So Nakagawa, Terumasa Ikeda and Kei Sato; 20fk0108451, to Takamasa Ueno, Akifumi Takaori-Kondo, G2P-Japan Consortium, Takashi Irie, Akatsuki Saito, So Nakagawa, Terumasa Ikeda and Kei Sato); AMED Research Program on HIV/AIDS (21fk0410034, to Akifumi Takaori-Kondo; 21fk0410033, to Akatsuki Saito; and 21fk0410039, to Kei Sato); AMED CRDF Global Grant (21jk0210039 to Akatsuki Saito); AMED Japan Program for Infectious Diseases Research and Infrastructure (21wm0325009, to Akatsuki Saito); JST A-STEP (JPMJTM20SL, to Terumasa Ikeda); JST SICORP (e-ASIA) (JPMJSC20U1, to Kei Sato); JST SICORP (JPMJSC21U5, to Kei Sato), JST CREST (JPMJCR20H6, to So Nakagawa; JPMJCR20H4, to Kei Sato); JSPS KAKENHI Grant-in-Aid for Scientific Research C (19K06382, to Akatsuki Saito); JSPS KAKENHI Grant-in-Aid for Scientific Research B (18H02662, to Kei Sato; and 21H02737, to Kei Sato); JSPS Fund for the Promotion of Joint International Research (Fostering Joint International Research) (18KK0447, to Kei Sato); JSPS Core-to-Core Program (A. Advanced Research Networks) (JPJSCCA20190008, to Kei Sato); JSPS Research Fellow DC1 (19J20488, to Izumi Kimura; xxxx, to Keiya Uriu); JSPS Leading Initiative for Excellent Young Researchers (LEADER) (to Terumasa Ikeda); The Tokyo Biochemical Research Foundation (to Kei Sato); Mitsubishi Foundation (to Terumasa Ikeda); Shin-Nihon Foundation of Advanced Medical Research (to Mako Toyoda and Terumasa Ikeda); Tsuchiya Foundation (to Takashi Irie); a Grant for Joint Research Projects of the Research Institute for Microbial Diseases, Osaka University (to Akatsuki Saito); an intramural grant from Kumamoto University COVID-19 Research Projects (AMABIE) (to Terumasa Ikeda); Intercontinental Research and Educational Platform Aiming for Eradication of HIV/AIDS (to Terumasa Ikeda); and Joint Usage/Research Center program of Institute for Frontier Life and Medical Sciences, Kyoto University (to Kei Sato).

## Consortia

### The Genotype to Phenotype Japan (G2P-Japan) Consortium

Shigeru Fujita^1^, Mai Suganami^1^, Akiko Oide^1^, Mika Chiba^1^, Naoko Misawa^1^, Takasuke Fukuhara^21^, Keita Matsuno^21^, Hirofumi Sawa^21^, Shinya Tanaka^21^, Tomokazu Tamura^21^, Rigel Suzuki^21^, Yuhei Morioka^21^, Kana Tsushima^21^, Haruko Kubo^21^, Naganori Nao^21^, Asako Shigeno^21^, Masumi Tsuda^21^, Mai Kishimoto^21^, Lei Wang^21^, Yoshitaka Oda^21^, Zannatul Ferdous^21^, Hiromi Mouri^21^, Miki Iida^21^, Keiko Kasahara^21^, Koshiro Tabata^21^, Mariko Ishizuka^21^, Kenzo Tokunaga^22^, Seiya Ozono^22^, Isao Yoshida^12^, Mami Nagashima^12^, Miyoko Takahashi^7^, Yasuhiro Kazuma^9^, Ryosuke Nomura^9^, Yoshihito Horisawa^9^, Yusuke Tashiro^9^, Yugo Kawai^9^, Ryoko Kawabata^13^, Otowa Takahashi^3^, Kimiko Ichihara^3^, Kazuko Kitazato^3^, Haruyo Hasebe^3^, Chihiro Motozono^11^, Isaac Ngare^11^

^21^ Hokkaido University, Sapporo, Japan.

^22^ National Institute of Infectious Diseases, Tokyo, Japan

## Methods

### Ethics statement

All experiments with hamsters were performed in accordance with the Science Council of Japan’s Guidelines for the Proper Conduct of Animal Experiments. The protocols were approved by the Institutional Animal Care and Use Committee of National University Corporation Hokkaido University (approval ID: 20-0060). All protocols involving specimens from human subjects recruited at Kyoto University and Kuramochi Clinic Interpark were reviewed and approved by the Institutional Review Boards of Kyoto University (approval ID: G1309) and Kuramochi Clinic Interpark (approval ID: G2021-004). All human subjects provided written informed consent. All protocols for the use of human specimens were reviewed and approved by the Institutional Review Boards of The Institute of Medical Science, The University of Tokyo (approval IDs: 2021-1-0416 and 2021-18-0617), Kyoto University (approval ID: G0697), Kumamoto University (approval IDs: 2066 and 2074), and University of Miyazaki (approval ID: O-1021).

### Human serum collection

Vaccine sera were collected from eleven vaccinees four weeks after their second vaccination with the BNT162b2 (Pfizer/BioNTech) vaccine (average age: 35, range: 29-56, 18% male) and sixteen vaccinees four weeks after their second mRNA-1273 (Moderna) vaccine (average age: 27, range: 20-47, 38% male).

Convalescent sera were collected from vaccine-naïve individuals who had been infected with the Delta variant (n=10; average age: 47, range: 22-63, 70% male). To identify the SARS-CoV-2 variants infecting patients, saliva was collected from COVID-19 patients during infection onset, and RNA was extracted using a QIAamp viral RNA mini kit (Qiagen, Cat# 52906) according to the manufacturer’s protocol. To identify the Delta variants, viral genome sequencing was performed as previously described (Meng *et al*., 2022). For details, see the “Viral genome sequencing” section below. Sera collected from twelve convalescents during the early pandemic (until May 2020) (average age: 71, range: 52-92, 8% male) were purchased from RayBiotech. Sera were inactivated at 56°C for 30 min and stored at –80°C until use. The details of the sera used in this study are summarized in **Table S1**.

### Hamster serum collection

Animal experiments were performed as previously described (Saito *et al*., 2022; Suzuki *et al*., 2022; Yamasoba *et al*., 2022). Syrian hamsters (male, 4 weeks old) were purchased from Japan SLC Inc. (Shizuoka, Japan). For the virus infection experiments, hamsters were anaesthetized by intramuscular injection of a mixture of either 0.15 mg/kg medetomidine hydrochloride (Domitor®, Nippon Zenyaku Kogyo), 2.0 mg/kg midazolam (FUJIFILM Wako Chemicals, Cat# 135-13791) and 2.5 mg/kg butorphanol (Vetorphale®, Meiji Seika Pharma), or 0.15 mg/kg medetomidine butorphanol. The chimeric recombinant SARS-CoV-2 (rB.1.1 S-GFP and rBA.1 S-GFP) (10,000 TCID_50_ in 100 μl) were intranasally inoculated under anesthesia (Yamasoba *et al*., 2022). Sera of infected hamsters were collected at 16 days postinfection (d.p.i.) using cardiac puncture under anesthesia with isoflurane and stored at −80°C until use.

### Cell culture

HEK293T cells (a human embryonic kidney cell line; ATCC, CRL-3216), HEK293 cells (a human embryonic kidney cell line; ATCC CRL-1573), and HOS-ACE2/TMPRSS2 cells, HOS cells stably expressing human ACE2 and TMPRSS2 (Ferreira et al., 2021; Ozono et al., 2021) were maintained in DMEM (high glucose) (Sigma-Aldrich, Cat# 6429-500ML) containing 10% fetal bovine serum (FBS) and 1% penicillin-streptomycin (PS). HEK293-C34 cells, *IFNAR1* KO HEK293 cells expressing human ACE2 and TMPRSS2 by doxycycline treatment (Torii *et al*., 2021), were maintained in Dulbecco’s modified Eagle’s medium (high glucose) (Sigma-Aldrich, Cat# R8758-500ML) containing 10% FBS, 10 μg/ml blasticidin (InvivoGen, Cat# ant-bl-1) and 1% PS.

VeroE6/TMPRSS2 cells (VeroE6 cells stably expressing human TMPRSS2; JCRB1819) (Matsuyama et al., 2020) were maintained in DMEM (low glucose) (Wako, Cat# 041-29775) containing 10% FBS, G418 (1 mg/ml; Nacalai Tesque, Cat# G8168-10ML) and 1% PS. Expi293F cells (Thermo Fisher Scientific, Cat# A14527) were maintained in Expi293 expression medium (Thermo Fisher Scientific, Cat# A1435101).

## METHOD DETAILS

### Viral genome sequencing

Viral genome sequencing was performed as previously described (Motozono *et al*., 2021; Saito *et al*., 2022; Suzuki *et al*., 2022; Yamasoba *et al*., 2022) with some modifications. Briefly, the virus sequences were verified by viral RNA-sequencing analysis. Viral RNA was extracted using a QIAamp viral RNA mini kit (Qiagen, Cat# 52906). The sequencing library employed for total RNA sequencing was prepared using the NEB Next Ultra RNA Library Prep Kit for Illumina (New England Biolabs, Cat# E7530). Paired-end 76-bp sequencing was performed using a MiSeq system (Illumina) with MiSeq reagent kit v3 (Illumina, Cat# MS-102-3001). Sequencing reads were trimmed using fastp v0.21.0 (Chen et al., 2018) and subsequently mapped to the viral genome sequences of a lineage A isolate (strain WK-521; GISAID ID: EPI_ISL_408667) (Matsuyama *et al*., 2020) using BWA-MEM v0.7.17 (Li and Durbin, 2009). Variant calling, filtering, and annotation were performed using SAMtools v1.9 (Li et al., 2009) and snpEff v5.0e (Cingolani et al., 2012).

### Molecular phylogenetic analyses

The SARS-CoV-2 genomes and annotation information used in this study were downloaded from the GISAID EpiCoV database (https://www.gisaid.org/) on January 8, 2022 (6,780,682 sequences). A total of 204,375 Omicron BA.1 variants were obtained, which included 1,074 B.1.1.529 variants because the B.1.1.529 lineage was recategorized as BA.1 as of February 24, 2022 (https://cov-lineages.org/lineage_list.html). For each sequence, we counted the number of undetermined nucleotides (such as N, Y, W) for whole genomes as well as *S* genes and obtained 40,739 sequences with fewer than 1,000 undetermined nucleotides in the genome and fewer than 10 undetermined nucleotides in the S-coding region. We then obtained BA.1 variant genomes that met the following criteria: 1) genomes were isolated from humans; 2) genomes did not contain any undetermined nucleotides in genomic regions corresponding to amino acid positions 371-375 in the S protein; 3) genomes were sampled from September 2021 to November 2021; and 4) genomes did not contain any of the 3 amino acid replacements in the S protein. We then selected 12 genomes and randomly selected 100 genomes that met criteria 1 and 2. By removing genomes that include possible recombination events by using RDP4 v4.101 (Martin et al., 2015), 48 Omicron genomes were obtained.

The 48 Omicron genomes with two outgroup genomes EPI_ISL_402125 (strain Wuhan-Hu-1, B lineage) and EPI_ISL_406862 (B.1 lineage; one of the earliest sequences carrying the S D614G mutation) were aligned using FFT-NS-1 in MAFFT suite v7.407 (Katoh and Standley, 2013). We then deleted gapped regions in the 5’ and 3’ regions. BEAST v1.10.4 (Suchard et al., 2018) was used to construct a timetree under an exponential growth coalescent model using a strict molecular clock. The GTR model with the four categories of discrete gamma rate variation was used as a nucleotide substitution model (Rodriguez et al., 1990; Yang, 1996). We ran Markov Chain Monte Carlo (MCMC) procedures with a 1 × 10^8^ chain length for all calculations, discarding the first 10% as burn-in and sampling every 10,000 replicates. The effective sample size for all run was confirmed to be larger than 200. FigTree v1.4.4 (http://tree.bio.ed.ac.uk/software/figtree/) was used to show the tree. To further determine the population history of the Omicron genomes, we generated a Bayesian skyline plot using the same model (2 × 10^8^ chain length for MCMC) and summarized the results using Tracer v1.7.1 (Rambaut et al., 2018).

### Plasmid construction

Plasmids expressing the codon-optimized SARS-CoV-2 S proteins of B.1 (the parental D614G-bearing variant), Delta (B.1.617.2) and Omicron (BA.1 lineage) variants were prepared in our previous studies (Ferreira *et al*., 2021; Kimura *et al*., 2022; Motozono *et al*., 2021; Saito *et al*., 2022; Suzuki *et al*., 2022; Uriu et al., 2022; Uriu et al., 2021; Yamasoba *et al*., 2022). Plasmids expressing a series of SAR-CoV-2 S mutants were generated by site-directed overlap extension PCR using the primers listed in **Table S3**. The resulting PCR fragment was digested with KpnI and NotI and inserted into the corresponding site of the pCAGGS vector (Niwa et al., 1991). Nucleotide sequences were determined by DNA sequencing services (Eurofins), and the sequence data were analyzed by Sequencher v5.1 software (Gene Codes Corporation).

### Pseudovirus assay

Pseudovirus assay was performed as previously described (Ferreira *et al*., 2021; Kimura *et al*., 2022; Motozono *et al*., 2021; Saito *et al*., 2022; Suzuki *et al*., 2022; Uriu *et al*., 2022; Uriu *et al*., 2021; Yamasoba *et al*., 2022). Briefly, lentivirus (HIV-1)-based, luciferase-expressing reporter viruses were pseudotyped with the SARS-CoV-2 spikes. HEK293T cells (500,000 cells) were cotransfected with 800 ng psPAX2-IN/HiBiT (Ozono et al., 2020), 800 ng pWPI-Luc2 (Ozono *et al*., 2020), and 400 ng plasmids expressing parental S or its derivatives using TransIT-293 Transfection Reagent (Takara, Cat# MIR2700) or or PEI Max (Polysciences, Cat# 24765-1) according to the manufacturer’s protocol. Two days posttransfection, the culture supernatants were harvested, and the pseudoviruses were stored at −80°C until use. The same amount of pseudoviruses (normalized to the HiBiT value, which indicates the amount of p24 HIV-1 antigen) was inoculated into HOS-ACE2/TMPRSS2. At two days postinfection, the infected cells were lysed with a Bright-Glo Luciferase Assay System (Promega, cat# E2620) or a One-Glo luciferase assay system (Promega, cat# E6130) and the luminescent signal was measured using a GloMax Explorer Multimode Microplate Reader (Promega) or a CentroXS3 plate reader (Berthhold Technologies).

### Western blot

Western blot was performed as previously described (Saito *et al*., 2022; Suzuki *et al*., 2022; Yamasoba *et al*., 2022). For the blot, the HEK293 cells cotransfected with the S expression plasmids and HIV-1-based pseudovirus producing plasmids (see “Pseudovirus assay” section above) or the HEK293 cells transfected with the S expression plasmids were used. To quantify the level of the cleaved S2 protein in the cells, the harvested cells were washed and lysed in lysis buffer [25 mM HEPES (pH 7.2), 20% glycerol, 125 mM NaCl, 1% Nonidet P40 substitute (Nacalai Tesque, Cat# 18558-54), protease inhibitor cocktail (Nacalai Tesque, Cat# 03969-21)]. After quantification of total protein by protein assay dye (Bio-Rad, Cat# 5000006), lysates were diluted with 2 × sample buffer [100 mM Tris-HCl (pH 6.8), 4% SDS, 12% β-mercaptoethanol, 20% glycerol, 0.05% bromophenol blue] and boiled for 10 m. Then, 10 μl samples (50 μg of total protein) were subjected to Western blot. To quantify the level of the S2 protein in the virions, 900 μl culture medium containing the pseudoviruses was layered onto 500 μl 20% sucrose in PBS and centrifuged at 20,000 g for 2 h at 4°C. Pelleted virions were resuspended in 1× NuPAGE LDS sample buffer (Thermo Fisher Scientific, Cat# NP0007) containing 2% β-mercaptoethanol and incubated at 70°C for 10 m. For protein detection, the following antibodies were used: mouse anti-SARS-CoV-2 S monoclonal antibody (clone 1A9, GeneTex, Cat# GTX632604, 1:10,000), mouse anti-HIV-1 p24 monoclonal antibody (183-H12-5C, obtained from the HIV Reagent Program, NIH, Cat# ARP-3537, 1:2,000), rabbit anti-beta actin (ACTB) monoclonal antibody (clone 13E5, Cell Signalling, Cat# 4970, 1:5,000), mouse anti-tubulin (TUBA) monoclonal antibody (clone DM1A, Sigma-Aldrich, Cat# T9026, 1:10,000), horseradish peroxidase (HRP)-conjugated horse anti-mouse IgG antibody (Cell Signaling, Cat# 7076S, 1:2,000), HRP-conjugated donkey anti-rabbit IgG polyclonal antibody (Jackson ImmunoResearch, Cat# 711-035-152, 1:10,000) and HRP-conjugated donkey anti-mouse IgG polyclonal antibody (Jackson ImmunoResearch, Cat# 715-035-150, 1:10,000). Chemiluminescence was detected using SuperSignal West Femto Maximum Sensitivity Substrate (Thermo Fisher Scientific, Cat# 34095), SuperSignal West Atto Ultimate Sensitivity Substrate (Thermo Fisher Scientific, Cat# A38554) or Western BLoT Ultra Sensitive HRP Substrate (Takara, Cat# T7104A) according to the manufacturer’s instruction. Bands were visualized using an Amersham Imager 600 (GE Healthcare) or iBright FL1500 Imaging System (Thermo Fisher Scientific), and the band intensity was quantified using Image Studio Lite v5.2 (LI-COR Biosciences) or Fiji software v2.2.0 (ImageJ).

### SARS-CoV-2 S-based fusion assay

SARS-CoV-2 S-based fusion assay was performed as previously described (Motozono *et al*., 2021; Saito *et al*., 2022; Suzuki *et al*., 2022; Yamasoba *et al*., 2022). This assay utilizes a dual split protein (DSP) encoding *Renilla* luciferase and *GFP* genes; the respective split proteins, DSP_8-11_ and DSP_1-7_, are expressed in effector and target cells by transfection. Briefly, on day 1, effector cells (i.e., S-expressing cells) and target cells (see below) were prepared at a density of 0.6–0.8 × 10^6^ cells in a 6-well plate. To prepare effector cells, HEK293 cells were cotransfected with the S expression plasmids (400 ng) and pDSP_8-11_ (400 ng) using TransIT-LT1 (Takara, Cat# MIR2300). To prepare target cells, HEK293 cells were cotransfected with pC-ACE2 (200 ng) and pDSP_1-7_ (400 ng). Target HEK293 cells in selected wells were cotransfected with pC-TMPRSS2 (40 ng) in addition to the plasmids above. HEK293-ACE2 cells and HEK293-ACE2/TMPRSS2 cells were transfected with pDSP_1-7_ (400ng). On day 3 (24 h posttransfection), 16,000 effector cells were detached and reseeded into 96-well black plates (PerkinElmer, Cat# 6005225), and target HEK293 cells were reseeded at a density of 1,000,000 cells/2 ml/well in 6-well plates. On day 4 (48 h posttransfection), target cells were incubated with EnduRen live cell substrate (Promega, Cat# E6481) at 37°C for 3 h and then detached, and 32,000 target cells were added to a 96-well plate with effector cells. *Renilla* luciferase activity was measured at the indicated time points using Centro XS3 LB960 (Berthhold Technologies). To measure the surface expression level of S protein, effector cells were stained with rabbit anti-SARS-CoV-2 S S1/S2 polyclonal antibody (Thermo Fisher Scientific, Cat# PA5-112048, 1:100). Normal rabbit IgG (SouthernBiotech, Cat# 0111-01, 1:100) was used as negative controls, and APC-conjugated goat anti-rabbit IgG polyclonal antibody (Jackson ImmunoResearch, Cat# 111-136-144, 1:50) was used as a secondary antibody. Surface expression level of S protein was measured using FACS Canto II (BD Biosciences) and the data were analyzed using FlowJo software v10.7.1 (BD Biosciences). To calculate fusion activity, *Renilla* luciferase activity was normalized to the MFI of surface S proteins. The normalized value (i.e., *Renilla* luciferase activity per the surface S MFI) is shown as fusion activity.

### SARS-CoV-2 reverse genetics

Recombinant SARS-CoV-2 was generated by circular polymerase extension reaction (CPER) as previously described (Motozono *et al*., 2021; Saito *et al*., 2022; Torii *et al*., 2021; Yamasoba *et al*., 2022). In brief, 9 DNA fragments encoding the partial genome of SARS-CoV-2 (strain WK-521, PANGO lineage A; GISAID ID: EPI_ISL_408667) (Matsuyama *et al*., 2020) were prepared by PCR using PrimeSTAR GXL DNA polymerase (Takara, Cat# R050A). A linker fragment encoding hepatitis delta virus ribozyme, bovine growth hormone poly A signal and cytomegalovirus promoter was also prepared by PCR. The corresponding SARS-CoV-2 genomic region and the PCR templates and primers used for this procedure are summarized in **Table S3**. The 10 obtained DNA fragments were mixed and used for CPER (Torii *et al*., 2021). To prepare GFP-expressing replication-competent recombinant SARS-CoV-2, we used fragment 9, in which the *GFP* gene was inserted in the *ORF7a* frame, instead of the authentic F9 fragment (**Table S3**) (Torii *et al*., 2021).

To generate chimeric recombinant SARS-CoV-2 (**Figures 1G and 5A**), mutations were inserted in fragment 8 by site-directed overlap extension PCR or the GENEART site-directed mutagenesis system (Thermo Fisher Scientific, Cat# A13312) according to the manufacturer’s protocol with the primers listed in **Table S3**. Recombinant SARS-CoV-2 that bears B.1 S [rB.1 S-GFP (virus I)] or Omicron S [rOmicron S-GFP (virus II)] was prepared in our previous studies (Saito *et al*., 2022; Yamasoba *et al*., 2022). Nucleotide sequences were determined by a DNA sequencing service (Fasmac), and the sequence data were analyzed by Sequencher v5.1 software (Gene Codes Corporation).

To produce chimeric recombinant SARS-CoV-2, the CPER products were transfected into HEK293-C34 cells using TransIT-LT1 (Takara, Cat# MIR2300) according to the manufacturer’s protocol. At 1 d posttransfection, the culture medium was replaced with Dulbecco’s modified Eagle’s medium (high glucose) (Sigma-Aldrich, Cat# R8758-500ML) containing 2% FCS, 1% PS and doxycycline (1 μg/ml; Takara, Cat# 1311N). At 7 d posttransfection, the culture medium was harvested and centrifuged, and the supernatants were collected as the seed virus. To remove the CPER products (i.e., SARS-CoV-2-related DNA), 1 ml of the seed virus was treated with 2 μl TURBO DNase (Thermo Fisher Scientific, Cat# AM2238) and incubated at 37°C for 1 h. Complete removal of the CPER products (i.e., SARS-CoV-2-related DNA) from the seed virus was verified by PCR. The working virus stock was prepared from the seed virus as described below (see “SARS-CoV-2 preparation and titration” section).

### SARS-CoV-2 preparation and titration

The working virus stocks of chimeric recombinant SARS-CoV-2 were prepared and titrated as previously described (Motozono *et al*., 2021; Saito *et al*., 2022; Suzuki *et al*., 2022; Torii *et al*., 2021; Yamasoba *et al*., 2022). In brief, 20 μl of the seed virus was inoculated into VeroE6/TMPRSS2 cells (5,000,000 cells in a T-75 flask). One hour post infection (h.p.i.), the culture medium was replaced with DMEM (low glucose) (Wako, Cat# 041-29775) containing 2% FBS and 1% PS. At 3 d.p.i., the culture medium was harvested and centrifuged, and the supernatants were collected as the working virus stock.

The titer of the prepared working virus was measured as the 50% tissue culture infectious dose (TCID_50_). Briefly, one day before infection, VeroE6/TMPRSS2 cells (10,000 cells) were seeded into a 96-well plate. Serially diluted virus stocks were inoculated into the cells and incubated at 37°C for 4 d. The cells were observed under microscopy to judge the CPE appearance. The value of TCID_50_/ml was calculated with the Reed–Muench method (Reed and Muench, 1938).

To verify the sequence of chimeric recombinant SARS-CoV-2, viral RNA was extracted from the working viruses using a QIAamp viral RNA mini kit (Qiagen, Cat# 52906) and viral genome sequence was analyzed as described above (see “Viral genome sequencing” section above). In brief, the viral sequences of *GFP*-encoding recombinant SARS-CoV-2 (strain WK-521; GISIAD ID: EPI_ISL_408667) (Matsuyama *et al*., 2020; Torii *et al*., 2021) that harbor the *S* genes of respective variants were used for the reference. Information on the unexpected mutations detected is summarized in **Table S4**, and the raw data are deposited in Gene Expression Omnibus (accession number: GSE196649 and xxxx).

### SARS-CoV-2 infection

SARS-CoV-2 infection was performed as previously described (Meng *et al*., 2022; Motozono *et al*., 2021; Saito *et al*., 2022; Suzuki *et al*., 2022; Yamasoba *et al*., 2022). Briefly, 1 d before infection, VeroE6/TMPRSS2 cells (10,000 cells) were seeded into a 96-well plate. SARS-CoV-2 (100 TCID_50_, m.o.i. 0.01) was inoculated and incubated at 37°C for 1 h. The infected cells were washed, and 180 μl of culture medium was added. The culture supernatant (10 μl) was harvested at the indicated timepoints and used for RT–qPCR to quantify the viral RNA copy number (see “RT–qPCR” section below).

### RT–qPCR

RT–qPCR was performed as previously described (Meng *et al*., 2022; Motozono *et al*., 2021; Saito *et al*., 2022; Suzuki *et al*., 2022; Yamasoba *et al*., 2022). Briefly, 5 μl of culture supernatant was mixed with 5 μl of 2 × RNA lysis buffer [2% Triton X-100, 50 mM KCl, 100 mM Tris-HCl (pH 7.4), 40% glycerol, 0.8 U/μl recombinant RNase inhibitor (Takara, Cat# 2313B)] and incubated at room temperature for 10 min. RNase-free water (90 μl) was added, and the diluted sample (2.5 μl) was used as the template for real-time RT-PCR performed according to the manufacturer’s protocol using the One Step TB Green PrimeScript PLUS RT-PCR kit (Takara, Cat# RR096A) and the following primers: Forward *N*, 5’-AGC CTC TTC TCG TTC CTC ATC AC-3’; and Reverse *N*, 5’-CCG CCA TTG CCA GCC ATT C-3’. The viral RNA copy number was standardized with a SARS-CoV-2 direct detection RT-qPCR kit (Takara, Cat# RC300A). Fluorescent signals were acquired using QuantStudio 3 Real-Time PCR system (Thermo Fisher Scientific), CFX Connect Real-Time PCR Detection system (Bio-Rad), Eco Real-Time PCR System (Illumina), qTOWER3 G Real-Time System (Analytik Jena) or 7500 Real-Time PCR System (Thermo Fisher Scientific).

### Fluorescence microscopy

Fluorescence microscopy was performed as previously described (Saito *et al*., 2022; Yamasoba *et al*., 2022). Briefly, 1 d before infection, VeroE6/TMPRSS2 cells (10,000 cells) were seeded into 96-well, glass bottom, black plates and infected with SARS-CoV-2 (100 TCID_50_, m.o.i. 0.01). At 24, 48, and 72 h.p.i., GFP fluorescence was observed under an All-in-One Fluorescence Microscope BZ-X800 (Keyence) in living cells, and a 13-square-millimeter area of each sample was scanned. under the same parameters. Images were reconstructed using an BZ-X800 analyzer software (Keyence), and the area and the fluorescent intensity of the GFP-positive cells was measured using this software.

### Plaque assay

Plaque assay was performed as previously described (Motozono *et al*., 2021; Saito *et al*., 2022; Suzuki *et al*., 2022; Yamasoba *et al*., 2022). Briefly, 1 d before infection, VeroE6/TMPRSS2 cells (100,000 cells) were seeded into a 24-well plate and infected with SARS-CoV-2 (1, 10, 100 and 1,000 TCID_50_) at 37°C for 2 h. Mounting solution containing 3% FBS and 1.5% carboxymethyl cellulose (Wako, Cat# 039-01335) was overlaid, followed by incubation at 37°C. At 3 d.p.i., the culture medium was removed, and the cells were washed with PBS three times and fixed with 4% paraformaldehyde phosphate (Nacalai Tesque, Cat# 09154-85). The fixed cells were washed with tap water, dried, and stained with staining solution [0.1% methylene blue (Nacalai Tesque, Cat# 22412-14) in water] for 30 m. The stained cells were washed with tap water and dried, and the size of plaques was measured using Adobe Photoshop 2021 v22.4.1 (Adobe).

### Neutralization assay

Neutralization assay was performed as previously described (Ferreira *et al*., 2021; Kimura *et al*., 2022; Mlcochova et al., 2021; Saito *et al*., 2022; Uriu *et al*., 2022; Uriu *et al*., 2021; Yamasoba *et al*., 2022). Briefly, pseudoviruses were prepared as described above (see “Pseudovirus assay” section). For the neutralization assay, the SARS-CoV-2 S pseudoviruses (counting ~20,000 relative light units) were incubated with serially diluted (40-fold or 120-fold to 29,160-fold dilution at the final concentration) heat-inactivated sera at 37°C for 1 h. Pseudoviruses without sera were included as controls. Then, an 80 μl mixture of pseudovirus and serum/antibody was added to HOS-ACE2/TMPRSS2 cells (10,000 cells/50 μl) in a 96-well white plate. At 2 d.p.i., pseudovirus infectivity was measured as described above (see “Pseudovirus assay” section). The assay of each serum was performed in triplicate, and the 50% neutralization titer (NT50) was calculated using Prism 9 (GraphPad Software).

### Protein structure

All protein structural analyses were performed using the PyMOL molecular graphics system v2.5.0 (Schrödinger). The crystal structures of SARS-CoV-2 D614G (B.1 lineage) S (PDB: 7KRQ) (Zhang *et al*., 2021) and Omicron S (PDB: 7T9J) (Mannar *et al*., 2022) were used. To predict inter-subunit interaction of the Omicron S trimer, each subunit of the D614G S trimer was replaced with the Omicron S monomer(Mannar *et al*., 2022). The distance between F375 and H505 was measured using the PyMOL molecular graphics system v2.5.0 (Schrödinger).

### Yeast surface display

Yeast surface display was performed as previously described (Dejnirattisai *et al*., 2022; Kimura *et al*., 2022; Motozono *et al*., 2021; Yamasoba *et al*., 2022; Zahradnik *et al*., 2021). Briefly, the carboxypeptidase domain of human ACE2 (residues 18-740) was expressed in Expi293F cells and purified by a 5-ml HisTrap Fast Flow column (Cytiva, Cat# 17-5255-01) and Superdex 200 16/600 (Cytiva, Cat# 28-9893-35) using an ÄKTA pure chromatography system (Cytiva), and the purified soluble ACE2 was labelled with CF640R (Biotium, Cat# 92108). Protein quality was verified using a Tycho NT.6 system (NanoTemper) and ACE2 activity assay kit (SensoLyte, Cat# AS-72086).

An enhanced yeast display platform for SARS-CoV-2 S RBD [wild-type (B.1.1), residues 336-528] yeast surface expression was established using *Saccharomyces cerevisiae* EBY100 strain and pJYDC1 plasmid (Addgene, Cat# 162458) as previously described (Dejnirattisai *et al*., 2022; Kimura *et al*., 2022; Motozono *et al*., 2021; Yamasoba *et al*., 2022; Zahradnik *et al*., 2021). To prepare a series of SARS-CoV-2 S RBD mutants, the site-directed mutagenesis was performed using the KAPA HiFi HotStart ReadyMix kit (Roche, Cat# KK2601) by restriction enzyme-free cloning procedure (Peleg and Unger, 2014). Primers for mutagenesis are listed in **Table S3**.

The binding affinities of SARS-CoV-2 S RBDs to human ACE2 were determined by flow cytometry titration experiments. The CF640R-labelled ACE2 at 12–14 different concentrations (200 nM to 13 pM in PBS supplemented with bovine serum albumin at 1 g/l) per measurement were incubated with expressed yeast aliquots and 10 nM bilirubin (Sigma-Aldrich, Cat# 14370-1G) and analyzed by using FACS S3e Cell Sorter device (Bio-Rad). The background binding subtracted fluorescent signal was fitted to a standard noncooperative Hill equation by nonlinear least-squares regression using Python v3.7 (https://www.python.org) as previously described (Zahradnik *et al*., 2021).

## QUANTIFICATION AND STATISTICAL ANALYSIS

In the single timepoint experiments, statistical significance was tested using a two-sided Student’s *t* test (**Figures 1B, 1E, 3B, 4B, 4E, 6D and 6E**), a two-sided paired *t* test (**Figures 1D, 4D, 6B and 6C**), a two-sided Mann–Whitney *U* test (**Figures 1J, 1K, 5D, 5E**), or a two-sided Wilcoxon signed-rank test (**Figure 2**). The tests above were performed using Prism 9 software v9.1.1 (GraphPad Software).

In the time-course experiments (**Figures 1F, 1H, 1I, 4F, 5B, 5C, 6F, and 6G**), a multiple regression analysis including experimental conditions (i.e., the types of infected viruses) as explanatory variables and timepoints as qualitative control variables was performed to evaluate the difference between experimental conditions thorough all timepoints. *P* value was calculated by a two-sided Wald test. Subsequently, familywise error rates (FWERs) were calculated by the Holm method. These analyses were performed in R v4.1.2 (https://www.r-project.org/).

In **Figures 1F, 1J, 6C, 6D and S1A**, assays were performed in triplicate. Photographs shown are the representatives of >18 fields of view taken for each sample.

---

**Figure S1.**
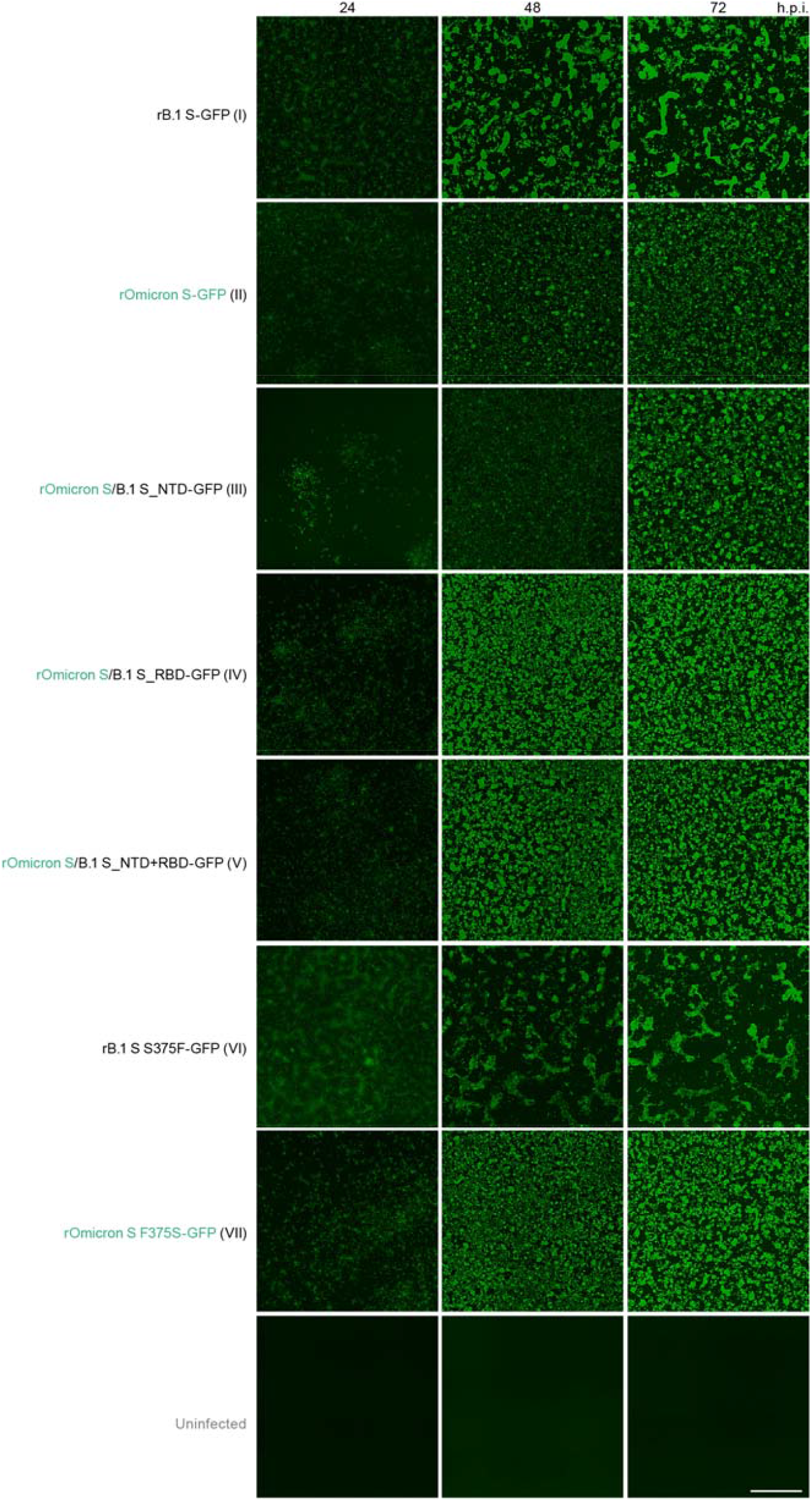
Representative images of chimeric recombinant SARS-CoV-2-infected cells, related to Figures 1 and 5. Fluorescence microscopy to measure the GFP area were measured in infected VeroE6/TMPRSS2 cells (m.o.i. 0.01) at 24, 48, and 72 h.p.i. The panels at 48 h.p.i. are identical to those shown in **Figures 1J and 5D**. Scale bar, 500 μm.

**Figure S2.**
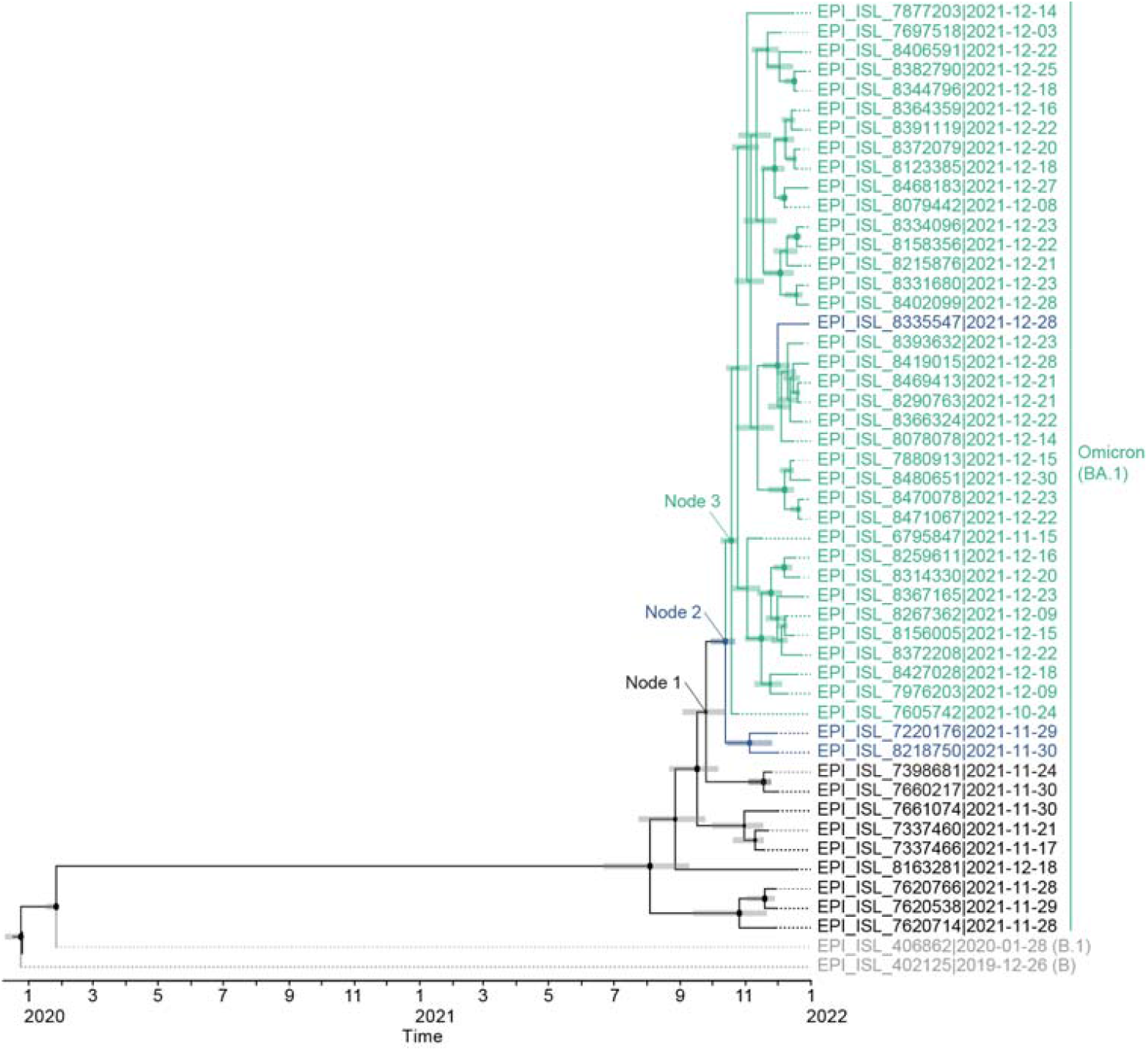
Evolutionary of Omicron. A timetree of 48 Omicron variants and two outgroups (B and B.1 lineages) with GISAID ID, PANGO lineage and sampling date. The topology of the phylogenetic tree is identical to that shown in **Figure 3C**, top. Green, Omicron variants containing S371L, S373P and S375F mutations; blue, Omicron variants containing the S371L and S373P mutations; black, Omicron variants without the S371L/S373P/S375F mutations; and gray, the two outgroups (B and B.1 lineages). Bars on each internal node indicate the 95% HPD interval of estimation time.

**Figure S3.**
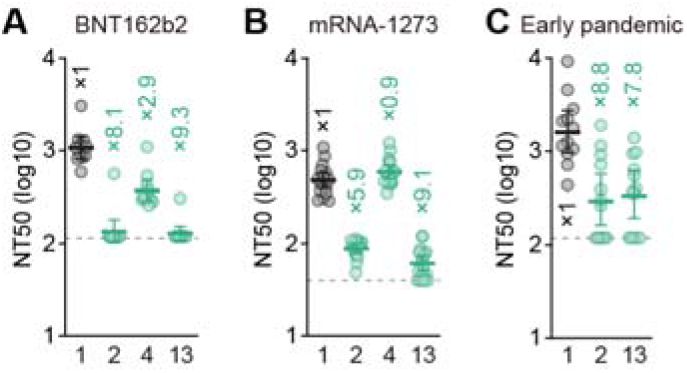
Immune resistance of the Omicron S S371L/S373P/S375F mutant. Neutralization assays were performed with pseudoviruses harboring a series of S proteins (summarized in **Figures 1A and 4A**). The numbers are identical to those in **Figures 1A and 4A**. Vaccinated sera [BNT162b2 (**A**, 11 donors); or mRNA-1273 (**B**, 16 donors)] and convalescent sera of individuals infected with an early pandemic virus (until May 2020) (**C**, 12 donors) were used. The list of sera used in this experiment is shown in **Table S1**. Each serum sample was tested in triplicate to determine the 50% neutralization titer (NT50). Each dot represents one NT50 value, and the geometric mean and 95% CI are shown. The numbers indicate the fold changes of resistance versus each antigenic variant. Horizontal gray lines indicate the detection limit of each assay (120 for **A and C**; 40 for **B**).

**Table S1. Human sera used in this study, related to Figure 2**

**Table S2. Mutations detected in the Omicron RBD, related to Figure 3.**

**Table S3. Primers used in this study, related to Figures 1 and 3–6.**

**Table S4. Summary of the mutations detected in the working virus stocks compared to the original sequences, related to Figures 1 and 5**

## References

Cameroni, E., Bowen, J.E., Rosen, L.E., Saliba, C., Zepeda, S.K., Culap, K., Pinto, D., VanBlargan, L.A., De Marco, A., di Iulio, J., et al. (2021). Broadly neutralizing antibodies overcome SARS-CoV-2 Omicron antigenic shift. Nature. 10.1038/s41586-021-04386-2.

Cao, Y., Wang, J., Jian, F., Xiao, T., Song, W., Yisimayi, A., Huang, W., Li, Q., Wang, P., An, R., et al. (2021). Omicron escapes the majority of existing SARS-CoV-2 neutralizing antibodies. Nature, doi: https://doi.org/10.1038/d41586-41021-03796-41586.

Carreño, J.M., Alshammary, H., Tcheou, J., Singh, G., Raskin, A., Kawabata, H., Sominsky, L., Clark, J., Adelsberg, D.C., Bielak, D., et al. (2021). Activity of convalescent and vaccine serum against SARS-CoV-2 Omicron. Nature, doi: https://doi.org/10.1038/d41586-41021-03846-z.

Cele, S., Jackson, L., Khoury, D.S., Khan, K., Moyo-Gwete, T., Tegally, H., San, J.E., Cromer, D., Scheepers, C., Amoako, D., et al. (2021). Omicron extensively but incompletely escapes Pfizer BNT162b2 neutralization. Nature, doi: https://doi.org/10.1038/d41586-41021-03824-41585.

Chen, S., Zhou, Y., Chen, Y., and Gu, J. (2018). fastp: an ultra-fast all-in-one FASTQ preprocessor. Bioinformatics 34, i884–i890. 10.1093/bioinformatics/bty560.

Cingolani, P., Platts, A., Wang le, L., Coon, M., Nguyen, T., Wang, L., Land, S.J., Lu, X., and Ruden, D.M. (2012). A program for annotating and predicting the effects of single nucleotide polymorphisms, SnpEff: SNPs in the genome of Drosophila melanogaster strain w1118; iso-2; iso-3. Fly (Austin) 6, 80–92. 10.4161/fly.19695.

Davies, N.G., Abbott, S., Barnard, R.C., Jarvis, C.I., Kucharski, A.J., Munday, J.D., Pearson, C.A.B., Russell, T.W., Tully, D.C., Washburne, A.D., et al. (2021). Estimated transmissibility and impact of SARS-CoV-2 lineage B.1.1.7 in England. Science 372. 10.1126/science.abg3055.

Dejnirattisai, W., Huo, J., Zhou, D., Zahradnik, J., Supasa, P., Liu, C., Duyvesteyn, H.M.E., Ginn, H.M., Mentzer, A.J., Tuekprakhon, A., et al. (2022). SARS-CoV-2 Omicron-B.1.1.529 leads to widespread escape from neutralizing antibody responses. Cell 185, 467–484 e415. 10.1016/j.cell.2021.12.046.

Dejnirattisai, W., Shaw, R.H., Supasa, P., Liu, C., Stuart, A.S., Pollard, A.J., Liu, X., Lambe, T., Crook, D., Stuart, D.I., et al. (2021). Reduced neutralisation of SARS-CoV-2 omicron B.1.1.529 variant by post-immunisation serum. Lancet, doi:https://doi.org/10.1016/S0140-6736(1021)02844-02840.

Ferreira, I., Kemp, S.A., Datir, R., Saito, A., Meng, B., Rakshit, P., Takaori-Kondo, A., Kosugi, Y., Uriu, K., Kimura, I., et al. (2021). SARS-CoV-2 B.1.617 mutations L452R and E484Q are not synergistic for antibody evasion. J Infect Dis 224, 989–994. 10.1093/infdis/jiab368.

Garcia-Beltran, W.F., Denis, K.J.S., Hoelzemer, A., Lam, E.C., Nitido, A.D., Sheehan, M.L., Berrios, C., Ofoman, O., Chang, C.C., Hauser, B.M., et al. (2021). mRNA-based COVID-19 vaccine boosters induce neutralizing immunity against SARS-CoV-2 Omicron variant. Cell, doi: https://doi.org/10.1016/j.cell.2021.1012.1033.

Hou, Y.J., Chiba, S., Halfmann, P., Ehre, C., Kuroda, M., Dinnon, K.H., 3rd, Leist, S.R., Schafer, A., Nakajima, N., Takahashi, K., et al. (2020). SARS-CoV-2 D614G variant exhibits efficient replication ex vivo and transmission in vivo. Science 370, 1464–1468. 10.1126/science.abe8499.

Katoh, K., and Standley, D.M. (2013). MAFFT multiple sequence alignment software version 7: improvements in performance and usability. Mol Biol Evol 30, 772–780. 10.1093/molbev/mst010.

Kimura, I., Kosugi, Y., Wu, J., Zahradnik, J., Yamasoba, D., Butlertanaka, E.P., Tanaka, Y.L., Uriu, K., Liu, Y., Morizako, N., et al. (2022). The SARS-CoV-2 Lambda variant exhibits enhanced infectivity and immune resistance. Cell Rep 38, 110218. 10.1016/j.celrep.2021.110218.

Korber, B., Fischer, W.M., Gnanakaran, S., Yoon, H., Theiler, J., Abfalterer, W., Hengartner, N., Giorgi, E.E., Bhattacharya, T., Foley, B., et al. (2020). Tracking changes in SARS-CoV-2 spike: evidence that D614G increases infectivity of the COVID-19 virus. Cell 182, 812–827. 10.1016/j.cell.2020.06.043.

Li, H., and Durbin, R. (2009). Fast and accurate short read alignment with Burrows-Wheeler transform. Bioinformatics 25, 1754–1760. 10.1093/bioinformatics/btp324.

Li, H., Handsaker, B., Wysoker, A., Fennell, T., Ruan, J., Homer, N., Marth, G., Abecasis, G., Durbin, R., and Genome Project Data Processing Subgroup (2009). The sequence alignment/map format and SAMtools. Bioinformatics 25, 2078–2079. 10.1093/bioinformatics/btp352.

Li, Q., Wu, J., Nie, J., Zhang, L., Hao, H., Liu, S., Zhao, C., Zhang, Q., Liu, H., Nie, L., et al. (2020). The impact of mutations in SARS-CoV-2 spike on viral infectivity and antigenicity. Cell 182, 1284–1294 e1289. 10.1016/j.cell.2020.07.012.

Liu, L., Iketani, S., Guo, Y., Chan, J.F.-W., Wang, M., Liu, L., Luo, Y., Chu, H., Huang, Y., Nair, M.S., et al. (2021). Striking antibody evasion manifested by the Omicron variant of SARS-CoV-2. Nature, doi: https://doi.org/10.1038/d41586-41021-03826-41583.

Mannar, D., Saville, J.W., Zhu, X., Srivastava, S.S., Berezuk, A.M., Tuttle, K.S., Marquez, A.C., Sekirov, I., and Subramaniam, S. (2022). SARS-CoV-2 Omicron variant: Antibody evasion and cryo-EM structure of spike protein-ACE2 complex. Science 375, 760–764. 10.1126/science.abn7760.

Martin, D.P., Murrell, B., Golden, M., Khoosal, A., and Muhire, B. (2015). RDP4: detection and analysis of recombination patterns in virus genomes. Virus Evol 1, vev003. 10.1093/ve/vev003.

Martineza, C.R., and Iverson, B.L. (2012). Rethinking the term “pi-stacking”. Chemical Science 3, 2191–2201.

Matsuyama, S., Nao, N., Shirato, K., Kawase, M., Saito, S., Takayama, I., Nagata, N., Sekizuka, T., Katoh, H., Kato, F., et al. (2020). Enhanced isolation of SARS-CoV-2 by TMPRSS2-expressing cells. Proc Natl Acad Sci U S A 117, 7001–7003. 10.1073/pnas.2002589117.

Meng, B., Abdullahi, A., Ferreira, I.A.T.M., Goonawardane, N., Saito, A., Kimura, I., Yamasoba, D., Gerber, P.P., Fatihi, S., Rathore, S., et al. (2022). Altered TMPRSS2 usage by SARS-CoV-2 Omicron impacts tropism and fusogenicity. Nature. 10.1038/s41586-022-04474-x.

Mlcochova, P., Kemp, S.A., Dhar, M.S., Papa, G., Meng, B., Ferreira, I., Datir, R., Collier, D.A., Albecka, A., Singh, S., et al. (2021). SARS-CoV-2 B.1.617.2 Delta variant replication and immune evasion. Nature 599, 114–119. 10.1038/s41586-021-03944-y.

Motozono, C., Toyoda, M., Zahradnik, J., Saito, A., Nasser, H., Tan, T.S., Ngare, I., Kimura, I., Uriu, K., Kosugi, Y., et al. (2021). SARS-CoV-2 spike L452R variant evades cellular immunity and increases infectivity. Cell Host Microbe 29, 1124–1136. 10.1016/j.chom.2021.06.006.

National Institute for Communicable Diseases, S.A. (2021). “New COVID-19 variant detected in South Africa (November 25, 2021)”. https://www.nicd.ac.za/new-covid-19-variant-detected-in-south-africa/.

Niwa, H., Yamamura, K., and Miyazaki, J. (1991). Efficient selection for high-expression transfectants with a novel eukaryotic vector. Gene 108, 193–199. 10.1016/0378-1119(91)90434-d.

Ozono, S., Zhang, Y., Ode, H., Sano, K., Tan, T.S., Imai, K., Miyoshi, K., Kishigami, S., Ueno, T., Iwatani, Y., et al. (2021). SARS-CoV-2 D614G spike mutation increases entry efficiency with enhanced ACE2-binding affinity. Nat Commun 12, 848. 10.1038/s41467-021-21118-2.

Ozono, S., Zhang, Y., Tobiume, M., Kishigami, S., and Tokunaga, K. (2020). Super-rapid quantitation of the production of HIV-1 harboring a luminescent peptide tag. J Biol Chem 295, 13023–13030. 10.1074/jbc.RA120.013887.

Peleg, Y., and Unger, T. (2014). Application of the restriction-free (RF) cloning for multicomponents assembly. Methods Mol Biol 1116, 73–87. 10.1007/978-1-62703-764-8_6.

Planas, D., Saunders, N., Maes, P., Guivel-Benhassine, F., Planchais, C., Buchrieser, J., Bolland, W.-H., Porrot, F., Staropoli, I., Lemoine, F., et al. (2021). Considerable escape of SARS-CoV-2 Omicron to antibody neutralization. Nature, doi: https://doi.org/10.1038/d41586-41021-03827-41582.

Plante, J.A., Liu, Y., Liu, J., Xia, H., Johnson, B.A., Lokugamage, K.G., Zhang, X., Muruato, A.E., Zou, J., Fontes-Garfias, C.R., et al. (2020). Spike mutation D614G alters SARS-CoV-2 fitness. Nature. 10.1038/s41586-020-2895-3.

Rambaut, A., Drummond, A.J., Xie, D., Baele, G., and Suchard, M.A. (2018). Posterior summarization in Bayesian phylogenetics using Tracer 1.7. Syst Biol 67, 901–904. 10.1093/sysbio/syy032.

Reed, L.J., and Muench, H. (1938). A simple method of estimating fifty percent endpoints. Am J Hygiene 27, 493–497.

Rodriguez, F., Oliver, J.L., Marin, A., and Medina, J.R. (1990). The general stochastic model of nucleotide substitution. J Theor Biol 142, 485–501. 10.1016/s0022-5193(05)80104-3.

Saito, A., Irie, T., Suzuki, R., Maemura, T., Nasser, H., Uriu, K., Kosugi, Y., Shirakawa, K., Sadamasu, K., Kimura, I., et al. (2022). Enhanced fusogenicity and pathogenicity of SARS-CoV-2 Delta P681R mutation. Nature 602, 300–306. 10.1038/s41586-021-04266-9.

Suchard, M.A., Lemey, P., Baele, G., Ayres, D.L., Drummond, A.J., and Rambaut, A. (2018). Bayesian phylogenetic and phylodynamic data integration using BEAST 1.10. Virus Evol 4, vey016. 10.1093/ve/vey016.

Suzuki, R., Yamasoba, D., Kimura, I., Wang, L., Kishimoto, M., Ito, J., Morioka, Y., Nao, N., Nasser, H., Uriu, K., et al. (2022). Attenuated fusogenicity and pathogenicity of SARS-CoV-2 Omicron variant. Nature. 10.1038/s41586-022-04462-1.

Takashita, E., Kinoshita, N., Yamayoshi, S., Sakai-Tagawa, Y., Fujisaki, S., Ito, M., Iwatsuki-Horimoto, K., Chiba, S., Halfmann, P., Nagai, H., et al. (2022). Efficacy of antibodies and antiviral drugs against Covid-19 Omicron variant. N Engl J Med. 10.1056/NEJMc2119407.

Torii, S., Ono, C., Suzuki, R., Morioka, Y., Anzai, I., Fauzyah, Y., Maeda, Y., Kamitani, W., Fukuhara, T., and Matsuura, Y. (2021). Establishment of a reverse genetics system for SARS-CoV-2 using circular polymerase extension reaction. Cell Rep 35, 109014.

Uriu, K., Cardenas, P., Munoz, E., Barragan, V., Kosugi, Y., Shirakawa, K., Takaori-Kondo, A., Sato, K., Ecuador-Covid19 Consortium, and The Genotype to Phenotype Japan (G2P-Japan) Consortium (2022). Characterization of the immune resistance of SARS-CoV-2 Mu variant and the robust immunity induced by Mu infection. J Infect Dis. 10.1093/infdis/jiac053.

Uriu, K., Kimura, I., Shirakawa, K., Takaori-Kondo, A., Nakada, T.A., Kaneda, A., Nakagawa, S., Sato, K., and The Genotype to Phenotype Japan (G2P-Japan) Consortium (2021). Neutralization of the SARS-CoV-2 Mu variant by convalescent and vaccine serum. N Engl J Med 385, 2397–2399. 10.1056/NEJMc2114706.

VanBlargan, L.A., Errico, J.M., Halfmann, P.J., Zost, S.J., Crowe, J.E., Jr., Purcell, L.A., Kawaoka, Y., Corti, D., Fremont, D.H., and Diamond, M.S. (2022). An infectious SARS-CoV-2 B.1.1.529 Omicron virus escapes neutralization by therapeutic monoclonal antibodies. Nat Med. 10.1038/s41591-021-01678-y.

WHO (2022). “Tracking SARS-CoV-2 variants (March 22, 2022)”. https://www.who.int/en/activities/tracking-SARS-CoV-2-variants/.

Wu, F., Zhao, S., Yu, B., Chen, Y.M., Wang, W., Song, Z.G., Hu, Y., Tao, Z.W., Tian, J.H., Pei, Y.Y., et al. (2020). A new coronavirus associated with human respiratory disease in China. Nature 579, 265–269. 10.1038/s41586-020-2008-3.

Yamasoba, D., Kimura, I., Nasser, H., Morioka, Y., Nao, N., Ito, J., Uriu, K., Tsuda, M., Zahradnik, J., Shirakawa, K., et al. (2022). Virological characteristics of SARS-CoV-2 BA.2 variant. BioRxiv, doi: https://doi.org/10.1101/2022.1102.1114.480335.

Yang, Z. (1996). Among-site rate variation and its impact on phylogenetic analyses. Trends Ecol Evol 11, 367–372. 10.1016/0169-5347(96)10041-0.

Yurkovetskiy, L., Wang, X., Pascal, K.E., Tomkins-Tinch, C., Nyalile, T.P., Wang, Y., Baum, A., Diehl, W.E., Dauphin, A., Carbone, C., et al. (2020). Structural and functional analysis of the D614G SARS-CoV-2 spike protein variant. Cell 183, 739–751 e738. 10.1016/j.cell.2020.09.032.

Zahradnik, J., Marciano, S., Shemesh, M., Zoler, E., Harari, D., Chiaravalli, J., Meyer, B., Rudich, Y., Li, C., Marton, I., et al. (2021). SARS-CoV-2 variant prediction and antiviral drug design are enabled by RBD in vitro evolution. Nat Microbiol 6, 1188–1198. 10.1038/s41564-021-00954-4.

Zhang, J., Cai, Y., Xiao, T., Lu, J., Peng, H., Sterling, S.M., Walsh, R.M., Jr., Rits-Volloch, S., Zhu, H., Woosley, A.N., et al. (2021). Structural impact on SARS-CoV-2 spike protein by D614G substitution. Science 372, 525–530. 10.1126/science.abf2303.

